# Incomplete functional divergence drives genetic synergy during floral development in tomato

**DOI:** 10.1101/2025.11.19.689068

**Authors:** Natalia Gaarslev, Ying Xu, Eléonore Lizé, Irene Julca, Nathasha Glover, Sebastian Soyk

## Abstract

Genetic synergy arises from interactions between functionally related genes that control complex traits in plants and animals. Synergy occurs when the combined effect of multiple genes exceeds the additive contribution from each individual gene. However, genetic mechanisms that drive and maintain synergy remain largely elusive. Here, we investigated synergistic interactions among *SEPALLATA* MADS-box genes during floral development in tomato. We discovered that synergy emerges from duplicated genes that partitioned functions to regulate inflorescence architecture and floral organ identity. Moreover, synergistic interactions are reflected in non-additive expression changes that coordinate successive developmental stages. Finally, we demonstrate that synergy occurs due to residual redundancy on a dose-sensitive module guiding floral identity. These results indicate that synergy emerges as a relic of redundancy when functional divergence remains incomplete due to gene dosage constraints. Our work provides insights into mechanisms through which gene families diverge to produce the substrate for biological innovations during evolution.

## INTRODUCTION

Combining gene mutations can lead to synergistic effects that exceed the expected cumulative effects of each mutation alone. As a result, synergistic interactions can catalyze major leaps in trait adaptation by enabling more-than-additive changes during evolution ^1^. Genetic synergy is frequently observed between mutations in genes that are part of networks controlling growth and development ^2^. A common source of synergy is based on functional redundancy that arises from gene duplication ^2^, which produces redundant paralogs that, in the absence of selective pressure for retention, can follow different evolutionary fates ^3^. One of the paralogs may accumulate beneficial mutations to acquire novel functions and neofunctionalize, while the other retains the ancestral function. Alternatively, both paralogs may accumulate mutations to partition the ancestral function and subfunctionalize. Finally, one of the paralogs may accumulate mutations and become a nonfunctional pseudogene. However, even ancient paralogs that functionally diverged are often still co-expressed and maintain some degree of redundancy ^3^.

In plants, genetic synergy is frequently observed during the development of flowers and flower-bearing shoots (inflorescences). Flowers develop when shoot apical meristems cease the production of vegetative organs and transition to reproductive development, giving rise to the different organs that constitute a flower. In tomato (*Solanum lycopersicum*) and other sympodial plants, each vegetative meristem (VM) matures to a transition meristem (TM) and terminates in a floral meristem (FM) ^4^, which produces the first flower of the multi-flowered inflorescence (**Figure 1A**). The second flower is produced by a new sympodial inflorescence meristem (SIM) that is initiated at the flank of the first floral meristem to again terminate in a floral meristem and flower. The reiterated process of meristem initiation and termination results in a multi-flowered inflorescence with flowers arranged in zigzag (**Figure 1B-D**). The timely control of meristem maturation towards reproductive fate ensures inflorescence development. By contrast, subtle shifts in transcriptional programs that coordinate meristem transitions alter flower production and inflorescence architecture in tomato and other species ^5–9^. In tomato, precocious expression programs result in premature termination of meristems and less complex inflorescences ^10,11^, while delayed expression programs are associated with the overproliferation of inflorescence meristems that introduce branchpoints in the inflorescence ^5,12,13^.

**Figure 1:**
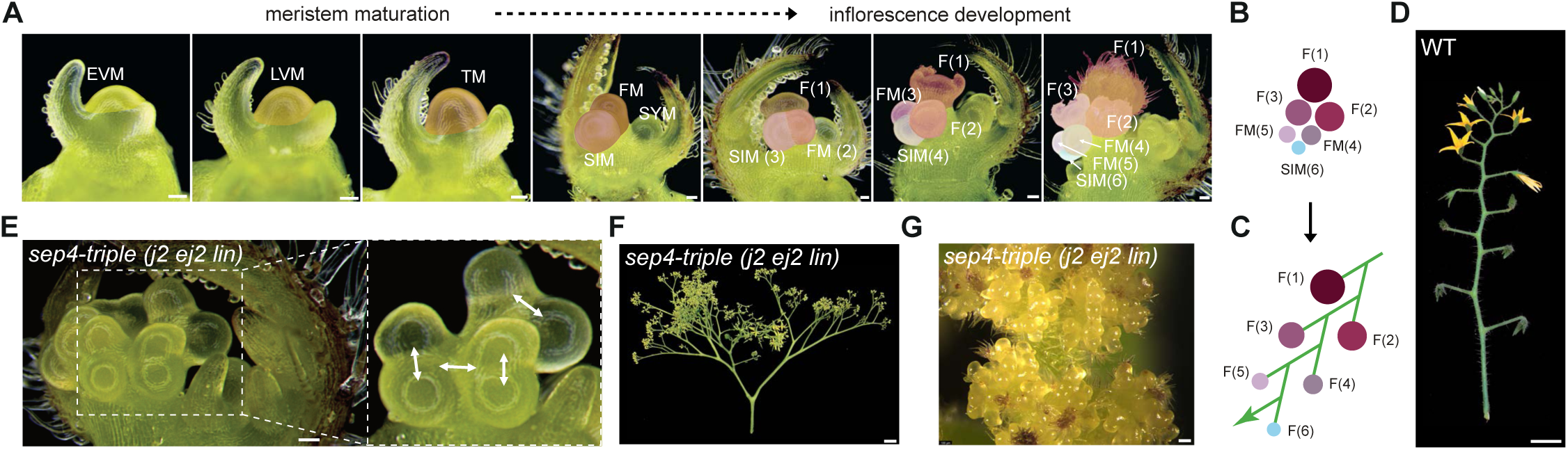
Synergy among tomato *SEP4* paralogs disturbs meristem maturation and leads to inflorescence branching. **(A)** Stereomicroscopy images of a wild-type (WT) shoot apex at subsequent stages of meristem maturation (EVM, early vegetative; LVM, late vegetative; TM, transition; FM, floral; SIM, sympodial inflorescence) and inflorescence development. SYM, sympodial vegetative meristem; F, flower. **(B)** Schematic representation of a developing inflorescence with three flowers, two FMs and one SIM arranged in the characteristic zig-zag pattern. **(C)** Diagram of a mature tomato inflorescence in which six subsequent flowers are arranged in a zig-zag pattern. The arrow represents a new SIM that is formed *de-novo* at the flank of the last flower. **(D)** Representative image of a wild-type tomato inflorescence (cv. Sweet-100). **(E)** Stereomicroscopy images of a *j2 ej2 lin* triple mutant shoot apex after the floral transition with additional inflorescence meristems. The orientation of the release of ectopic meristems is indicated by white arrows in the zoomed image. **(F)** Detached inflorescence of the *j2 ej2 lin* triple mutant showing excessive inflorescence branching and overproliferation of meristematic tissue leading to a cauliflower inflorescence phenotype. **(G)** Stereoscopy image of an *j2 ej2 lin* inflorescence. Scale bars indicate 100 µm (A, E, G) and 1cm (D, F).

Key regulators of meristem maturation have been identified in tomato. Mutations in *FALSIFLORA* (*FA*) ^14,15^, the ortholog of Arabidopsis *LEAFY* (*LFY*) ^16^, the *S/WUSCHEL RELATED HOMEOBOX9* (*S*/*SlWOX9*)^17^, and the F-box gene *ANANTHA* (*AN*) ^17,18^, ortholog of Arabidopsis *UNUSUAL FLORAL ORGANS* (*UFO*) ^19^ lead to the development of highly branched inflorescence structures that overproliferate meristems and resemble cauliflower curds. Moreover, the three *SEPALLATA* (*SEP*)-clade MADS-box (*MINICHROMOSOME MAINTENANCE1* (*MCM1*), *AGAMOUS* (*AG*), *DEFICIENS* (*DEF*), *SERUM RESPONSE FACTOR* (*SRF*)) transcription factor paralogs *JOINTLESS2* (*J2*), *ENHANCER OF J2* (*EJ2*) and *LONG INFLORESCENCE* (*LIN*) regulate the transition of meristems to floral identity and inflorescence branching patterns ^12^. Single *j2*, *ej2*, and *lin* mutants develop mainly unbranched inflorescences, while the *j2 ej2 lin* triple mutants bear cauliflower-like inflorescences that overproduce undifferentiated meristem tissue instead of flowers. The *j2 ej2 lin* triple mutant phenocopies the tomato *an* mutant and shows similarities to the Arabidopsis *apetala1 cauliflower* (*ap1 cal*) double mutants ^20–22^ (**Figure 1E-G**).

*SEP* functions were first reported in the context of floral organ specification ^23–25^. Floral organs arise from floral meristems, which typically produce organ primordia that are organized in concentric whorls, with outer vegetative organs, sepals and petals, protecting the inner reproductive organs, the stamens and carpels. Seminal work in Arabidopsis demonstrated that floral organ development is driven by homeotic genes classified into A-, B-, and C-classes ^26–28^. In this ABC model, all components, except *APETALA2* (*AP2*), encode MADS-box transcription factors. The A-class genes *AP1* and *AP2* specify sepals while the combination of A-class with the B-class genes *AP3* and *PISTILLATA* (*PI*) specifies petal identity. B-class and the C-class gene *AGAMOUS* (*AG*) determine the identity of stamens, while C-class function determines carpel identity and promotes floral meristem termination. In the early 2000s, the ABC model was expanded by the E-class function, which is required for proper organ development in all whorls and conferred by the *SEP* MADS-box clade with four members in Arabidopsis ^24,25^. At the molecular level, SEP proteins form tetrameric complexes with ABC-class MADS-box proteins in a specific spatiotemporal context to specify distinct floral organ identities ^29,30^. MADS-box tetramers consist of two homo- or heterodimers that bind to CArG-box DNA motifs and facilitate DNA looping between distant regulatory elements and target promoters to drive spatiotemporal expression patterns ^31^.

In Arabidopsis, *SEP* genes act largely redundantly during floral organ development. While *sep3* single mutants show subtle floral organ defects ^31^, *sep1234* quadruple mutants develop flowers entirely composed of organs with leaf-like identity ^24,25^. Studies in other species demonstrated functional divergence among *SEP* paralogs that emerged from *SEP4* duplications ^12,32,33^. Tomato encodes four *SEP4* paralogs, *J2*, *EJ2*, *LIN* and *RIPENING INHIBITOR* (*RIN*) ^12^. *J2* regulates the development of the fruit abscission zone while *EJ2* specifies sepal length and *LIN* regulates the number of flowers per inflorescence ^12,34,35^. The fourth *SEP4* paralog, *RIN*, is specifically expressed during fruit maturation and *rin* mutations delay fruit ripening ^36^ (**Figure S1A**). Genetic interactions between *J2, EJ2*, and *LIN* lead to synergistic effects on the rate of meristem maturation and inflorescence architecture in higher order mutants (**Figure 1E-G**) ^12^. A similar scenario has been described for *SEP4* paralogs in petunia, where the combined loss of *FLORAL BINDING PROTEIN4 (FBP4), FBP9,* and *FBP23* leads to highly branched inflorescences that fail to form flowers ^32^. Yet, the genetic and molecular mechanisms that drive synergy between *SEP* paralogs and consequences on gene expression programs underlying inflorescence development remain elusive.

## RESULTS

### Unequal redundancy between *SEP4* paralogs determines tomato inflorescence architecture

Combining mutations in the tomato *SEP4* paralogs *J2*, *EJ2* and *LIN* results in synergistic increases in inflorescence branching and transforms *j2 ej2 lin* triple mutant inflorescences into highly branched, cauliflower-like structures ^12^ (**Figure 1E-G**). To further dissect this example of genetic synergy into the individual contributions of *J2, EJ2,* and *LIN*, we analyzed single and double loss-of function mutants in a cherry tomato cultivar (cv. Sweet-100) for changes in meristem development and inflorescence branching (**Figure S1B**). Confirming previous findings, the *j2*, *ej2*, and *lin* single mutants were not markedly affected in inflorescence branching but exhibited specific single mutant phenotypes (**Figure S1C-F**). Fruit abscission was lost in *j2*, sepals were elongated in *ej2*, and inflorescences developed additional flowers in *lin* ^12^. In addition, *j2 ej2* double mutants produced ectopic inflorescence meristems, which resulted in strongly branched inflorescences on primary and sympodial shoots (**Figure 2A-C**). In comparison, *lin j2* and *lin ej2* double mutants initiated fewer ectopic inflorescence meristems, which led to weakly branched inflorescences at high penetrance on sympodial shoots and rarely on primary shoots (**Figure 2C-G**). The *lin j2* and *lin ej2* double mutants also developed more flowers per inflorescence compared to WT, but were not markedly enhanced compared to *lin* single mutants (**Figure 2H**). These observations indicate synergistic interactions between *LIN*/*J2* and *LIN*/*EJ2* during inflorescence development, however, the effect was weaker when compared with synergy between *J2/EJ2* ^12^. To account for genotype dependencies, we also revisited a previously published mutant collection in a plum tomato (cv. M82) ^12^, which confirmed weak inflorescence branching and more flowers per inflorescence in *lin j2* and *lin ej2* (**Figure 2I-J** and **Figure S1G**). In conclusion, we resolved synergistic interactions in *lin j2* and *lin ej2* double mutants that are weaker compared with interactions in *j2 ej2.* These results indicate unequal redundancy between *SEP4* paralogs with a minor contribution of *LIN* to the suppression of inflorescence branching.

**Figure 2:**
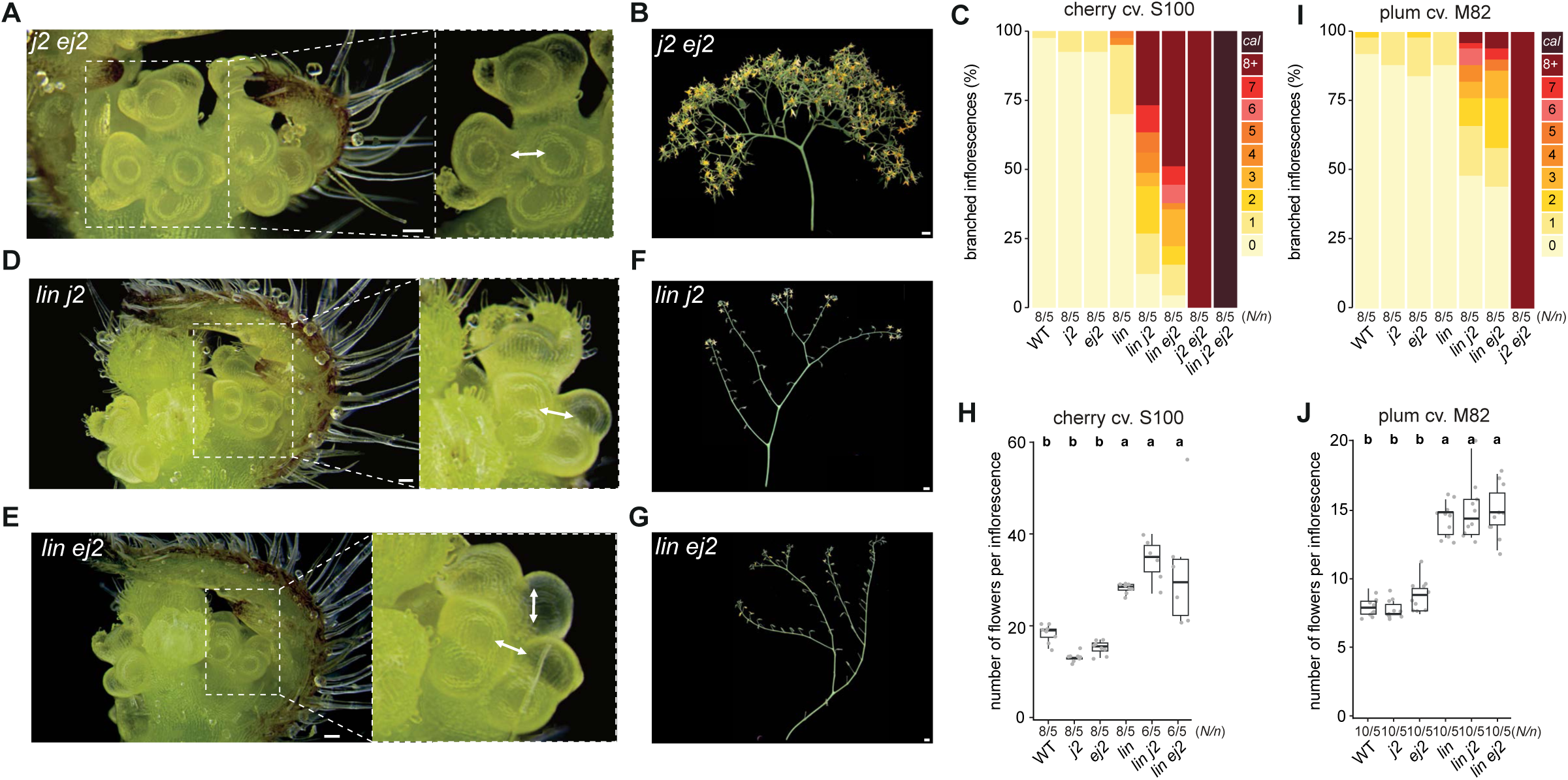
Unequal redundancy among the *SEP4* paralogs *J2*, *EJ2*, and *LIN* suppresses inflorescence branching and flower production in tomato. **(A)** Stereomicroscopy images of a *j2 ej2* double mutant shoot apex after the floral transition with additional ectopic inflorescence meristems on the primary inflorescence. The orientation of *de-novo* release of meristems on the primary inflorescence is indicated by white arrows in the zoomed image. Scale=100 µm. **(B)** Detached inflorescence of the *j2 ej2* double mutant (cv. S100) displaying strong inflorescence branching. Scale=1cm. **(C)** Quantification of inflorescence branching in the S100 cultivar shown as the percentage of branched inflorescences per genotype. cal, cauliflower inflorescence. **(D-E)** Stereoscope images of the shoot apex of *lin j2* (D) and *lin ej2* (E) double mutants. The orientation of the release of ectopic meristems on the inflorescence of the first sympodial shoot is indicated by white arrows in the zoomed image. Scale=100 µm. **(F-G)** Images of detached inflorescences from the *lin j2* (F) and *lin ej2* (G) double mutants showing weak branching. Scale=1cm. **(H)** Quantification of flower number per inflorescence in the S100 cultivar. Note that the *j2 ej2* mutant was excluded due to the unquantifiable (high) flower number. **(I)** Quantification of inflorescence branching in the M82 cultivar as in (C). **(J)** Quantification of flower number per inflorescence in the M82 cultivar as in (H). *N*/*n* in (C), (H-J) represent the number of plants (*N*) and inflorescences per plant (*n*) used for phenotyping. Letters in (H, J) represent the results from pairwise comparisons of means using one-way ANOVA and post hoc Tukey’s HSD test with 95% confidence level.

### *SEP4* synergy acts through a limited set of genes after meristems transition to floral fate

Inflorescence architecture is determined by the rate at which meristems mature from vegetative to reproductive identity ^5,13^. To gain molecular insights into how *SEP4* synergy affects the rate of meristem maturation, we sequenced mRNA from micro-dissected meristems of the single and double *sep4* mutants. We sampled meristems at the transition stage of meristem maturation (hereafter transition meristem samples), as well as a combined floral meristem/sympodial inflorescence meristem stage (hereafter floral meristem samples), during which *J2*, *EJ2*, and *LIN* are highly expressed (**Figure 1A** and **S1A**). We excluded the *j2 ej2 lin* triple mutant from this experiment due to the difficulty of obtaining meristem samples from a segregating triple mutant population. Multidimensional scaling (MDS) analyses of transcriptomes revealed that floral meristem samples clustered by the extent of inflorescence branching (**Figure 3A, B** and **Figure S2**) (see **Methods**). In contrast, transition meristem samples lacked clear clustering patterns, suggesting that floral meristem samples better captured relevant transcriptional changes associated with inflorescence branching. A differential expression analysis contrasting mutant genotypes with the WT yielded a total of 946 and 849 differentially expressed genes (DEGs; |log_2_FC| > 0.58, FDR<0.01) in the transition and floral meristem samples, respectively (**Figure 3C,D** and **Tables S1-S6**). A rather small fraction of DEGs was shared between the three double mutants in the transition samples (68 DEGs, 7.2%) (**Figure 3E** and **Table S7**), but this fraction increased in floral samples (104 DEGs, 12.3%) (**Figure 3F** and **Table S8**). Furthermore, while gene ontology (GO) enrichment analysis of the 946 DEGs from the transition meristem samples did not yield any significantly enriched terms, the 849 DEGs in floral meristem samples showed significant enrichment for GO categories related to morphogenesis, flower development, and floral organ identity (**Figure 3G-H**). We concluded that *SEP4* paralogs retained redundant functions to regulate a defined genetic module during inflorescence and floral meristem maturation stages, at which *SEP4* genes are highly expressed (**Figure S1A**). Combined loss of *SEP4* activity changes this genetic module and disturbs meristem maturation during the acquisition of floral identity.

**Figure 3:**
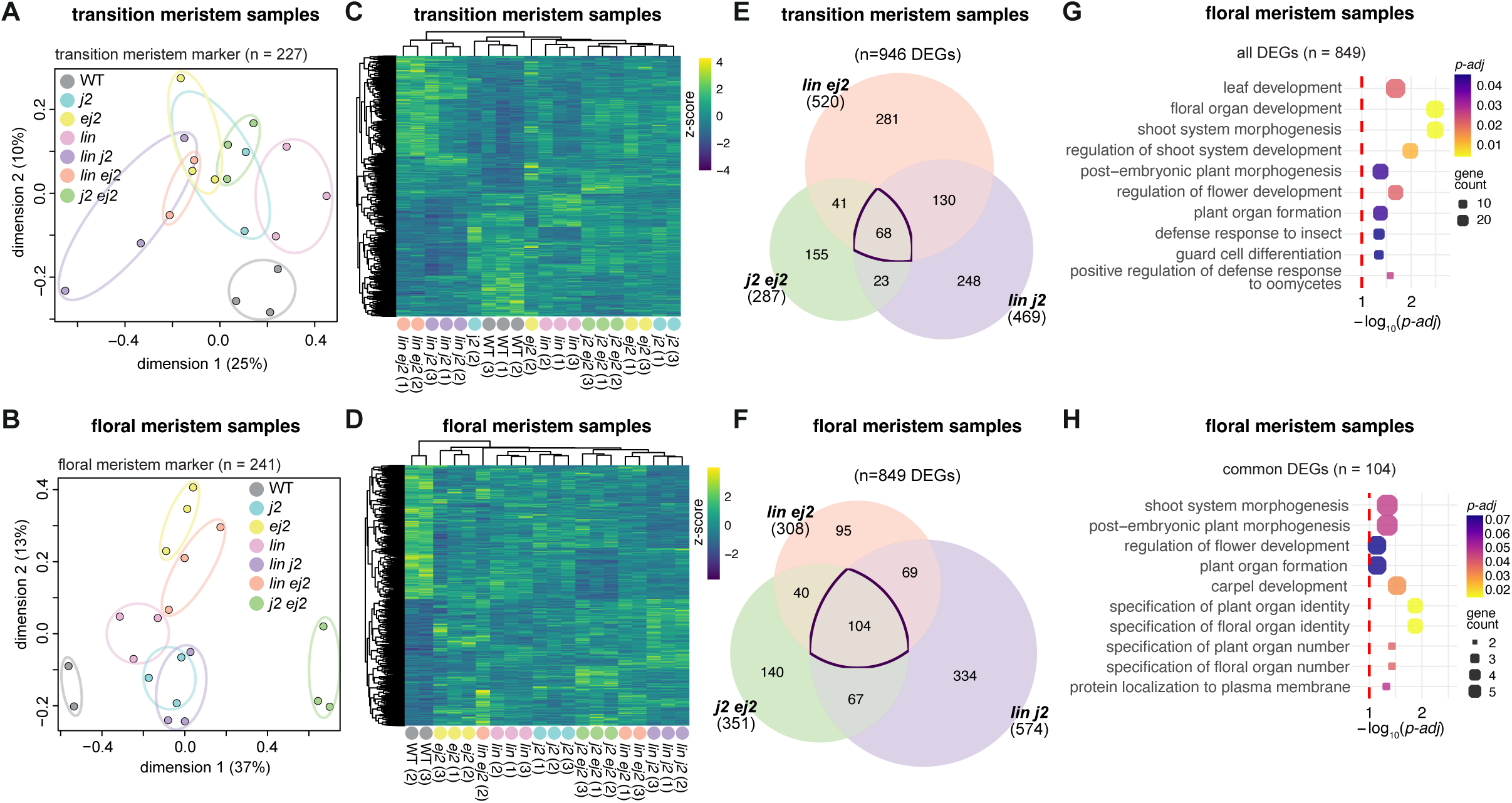
*SEP4* synergy acts on a shared module of genes after the transition stage of meristem maturation. **(A-B)** Multidimensional scaling (MDS) plots of gene expression of 227 transition marker genes in transition meristem samples (A) and 241 floral marker genes in floral meristem samples (B). **(C-D)** Heatmap showing z-score normalised expression of differentially expressed (FC ≥ 1.5, FDR ≤ 0.01) genes in transition (n=946) (C) and floral (n=849) (D) meristem samples. **(E-F)** Overlap of genes differentially expressed in transition (E) and floral (F) meristem samples in *j2 ej2*, *lin j2*, and *lin ej2* double mutants compared with the WT. The total number of DEGs per genotype is indicated in parenthesis. **(G-H)** The 10 most enriched gene ontology (GO) categories for all 849 (G) and common 104 (H) genes differentially expressed in *j2 ej2*, *lin j2*, and *lin ej2* floral meristem samples. Dot size is proportionate to the count of genes per term. *P* values obtained by Benjamini-Hochberg (BH) method in clusterProfiler.

### *SEP4* synergy is mirrored by a small group of genes with synergistic expression patterns

To pinpoint genetic modules that underlie synergistic changes in inflorescence branching, we focused on genes following synergistic expression patterns across floral meristem samples from single and double *sep4* mutants. We adapted an established framework ^37^ to compare relative expression changes in single and respective double mutants, and isolate genes with non-additive expression patterns that deviate from expected additive effects (**Figure 4A**, **Tables S1-S6**). By including directionality of non-additive expression changes relative to the predicted additive effects, we classified genes into three categories: (1) additive, in which the observed expression change in the double mutant did not significantly differ from the cumulative observed change in the two single mutants; (2) less-than-additive, in which the double mutant change was significantly smaller than the cumulative single mutant change; and (3) synergistic, in which the double mutant change was significantly greater than the cumulative single mutant change. Most genes followed non-additive (less-than-additive or synergistic) expression patterns (**Figure 4B-C**). For example, among the 351 DEGs in the *j2 ej2* floral meristem samples, 220 (63%) followed non-additive expression patterns, including 139 less-than-additive and 81 synergistic genes. Among the non-additive expression changes, we observed more less-than-additive than synergistic changes. DEGs with less-than-additive expression are likely controlled by MADS-box heterocomplexes that contain multiple SEP4 paralogs. In such cases, loss of a single *SEP4* paralog could already cause a reduction in heterocomplex activity that is not further enhanced in the double mutant. On this basis, less-than-additive DEGs unlikely explain the synergistic inflorescence phenotypes that are only observed in the double mutants. We concluded that synergistic inflorescence phenotypes are mirrored by widespread non-additive expression changes but only a small set of genes with synergistic changes drive these complex expression changes.

**Figure 4:**
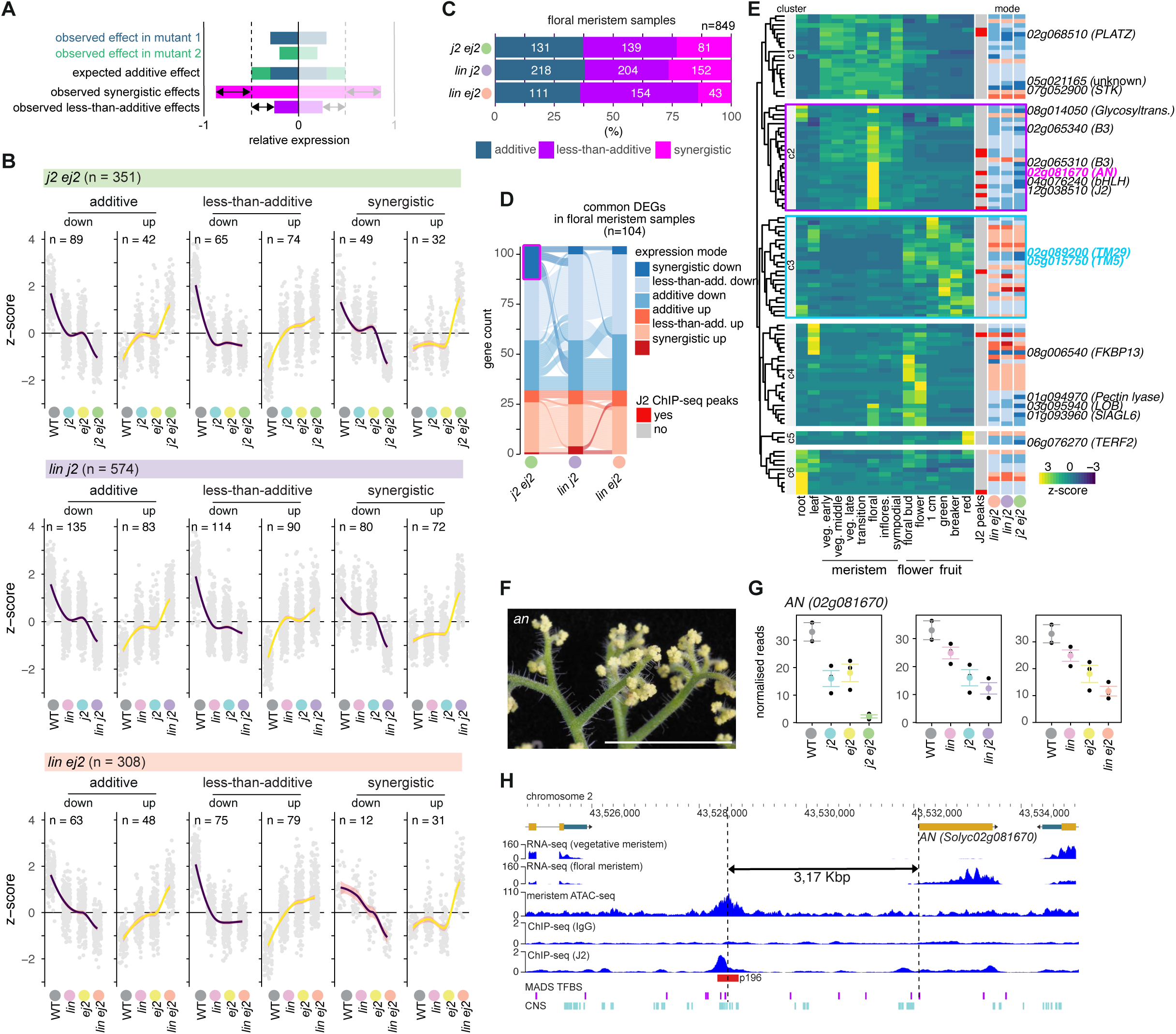
Synergy among *SEP4* paralogs acts on *ANANTHA* to drive the maturation of meristems towards floral fate. **(A)** Schematic representation of additive and non-additive gene expression categories in double mutants when compared with single mutants. **(B)** Normalized expression (z-score) of differentially expressed genes in floral meristems *j2 ej2* (top), *lin j2* (middle), and *lin ej2* (bottom) mutants compared with the WT, classified into additive, less-than-additive, and synergistic (more-than-additive) gene expression categories. Each category is split into genes that are down- and upregulated compared with the WT. Loess-smoothed trendlines are plotted in purple and yellow for down- and upregulated categories, respectively, with 95% confidence intervals shaded in red. n, number of genes per category. (C) Distribution of differentially expressed genes in *j2 ej2* (top), *lin j2* (middle), and *lin ej2* (bottom) mutants into additive, less-than-additive, and synergistic (more-than-additive) gene expression categories. **(D)** Overlap of gene expression categories in 104 gene differentially expressed in *j2 ej2*, *lin j2*, and *lin ej2*. **(E)** Normalized gene expression (z-score) for genes in (D) in different tissues and developmental stages. Both expression mode and presence of J2 ChIP-Seq peak are indicated as in (D). Genes synergistically downregulated in *j2 ej2* compared to the single mutants and WT are labelled with gene names. Magenta and blue boxes indicate gene clusters c2 (floral meristem) and c3 (flowers and fruit), respectively. **(F)** Macroscopic image of an *an* mutant inflorescence with overproliferating meristem tissue (cauliflower) instead of flowers. Scale = 1 cm. **(G)** Normalized gene expression of *AN* in floral meristem samples from single and double mutant combinations for *j2 ej2*, *lin j2*, and *lin ej2*. Data are represented as mean ± SE. **(H)** Browser view of the *AN* genomic region. Normalized coverage in counts per million (CPM) is shown for vegetative and floral meristem RNA-seq, meristem ATAC-Seq, and J2 ChIP-Seq (including IgG control). A significant J2 binding peak (P196) is indicated by a red square. Predicted MADS-box transcription factor binding sites (MADS TFBS) and conserved noncoding sequences (CNS) are indicted by magenta and cyan squares, respectively. Genomic positions on chromosome 2 (SL4.0) and the distance between the J2 binding site and the transcriptional start site (TSS) are indicated.

### *SEP4* synergy acts on the conserved floral meristem identity gene *ANANTHA*

To further investigate synergistic gene expression patterns that are associated with changes in inflorescence branching, we focused on the 104 DEGs that were shared between double mutants in floral meristem samples (**Figure 4D** and **Table S8**). Among those shared DEGs, the *j2 ej2* double mutant yielded the largest number of synergistic DEGs, consistent with the strong, synergistic inflorescence phenotype (**Figure 2B**). Most synergistic DEGs were downregulated in the *j2 ej2* mutant but downregulated with additive or less-than-additive patterns in the *lin j2* and *lin ej2* combinations, in agreement with the milder inflorescence phenotypes (**Figure 2F,G**). When we investigated tissue-specific expression patterns of 104 shared DEGs using public expression data, we identified a cluster of genes with peak expression in floral meristems ^5,38^ (**Figure 4E**). This floral meristem cluster (c2) included *AN*, a known regulator of floral meristem identity and ortholog of Arabidopsis *UFO* ^17,39^. Knockout of *AN* leads to cauliflower-like inflorescences reminiscent of the *j2 ej2 lin* triple mutants (**Figure 4F**) ^17,40^. *AN* expression was synergistically downregulated in *j2 ej2* but additively downregulated in *lin j2* and *lin ej2* genotypes, suggesting that *SEP4* dosage-dependent changes in *AN* expression underlie gradual changes in inflorescence branching (**Figure 4G**). This model is further supported by a J2 binding site upstream of *AN* that we identified in published Chromatin Immunoprecipitation-sequencing (ChIP-seq) data ^41^. Interestingly, the J2 binding site resides in an accessible chromatin region (ACR), which contains conserved noncoding sequences (CNS) and *cis*-regulatory elements (CREs) that were shown to impact *AN* transcription^42^ (**Figure 4H**). Given that J2 and other SEP factors bind to a conserved MADS-box target motif (CArG-box) ^41,43^, we propose *AN* as a direct transcriptional target of SEP4 factors. In conclusion, reductions in the dosage of *SEP4* paralogs result in quantitative downregulation of *AN*, which explains the synergistic effect on inflorescence branching. However, other genes with peak expression in floral meristems and similar synergistic expression patterns likely act as additional drivers of *SEP4* synergy.

### *SEP4* synergy is relayed to *SEP1/2/3* paralog expression levels

In addition to the floral meristem cluster (c2), we identified a cluster (c3) of genes that were primarily expressed in samples derived from developing flowers and fruits, and that included two additional *SEP* paralogs, *TOMATO MADS-BOX5* (*TM5*) and *TM29* (**Figure 4E**). Previous studies had shown that individual knock-down of *TM5* and *TM29* by antisense technology transforms carpels, the female reproductive floral organs that give rise to the tomato fruit, into ectopic inflorescences ^44,45^. *TM5* and *TM29* are lowly expressed during the floral meristem stage of meristem maturation (**Figure 4E**) and differentially expressed in floral meristem samples across all tomato *sep4* double mutants (**Figure 5A**), which suggested an additional layer of *SEP* paralog redundancy in floral meristems. Therefore, we revisited the phylogenetic relationship of tomato SEP proteins using hierarchical orthologous groups (HOGs) ^46^ (**Figure 5B**, **Figure S3A**, **Table S9-S10**). As previously reported ^12^, we detected an expansion of the SEP family in tomato to six members compared with only four in Arabidopsis (**Figure 5B-C**)^11^. A similar expansion had been observed in the SEP family of the related *Solanaceae* species petunia ^32^, and we observed more than four SEP proteins in all other *Solanaceae* species in our dataset (**Figure 5C**). This family expansion in *Solanaceae* is driven by duplication events in the SEP4 clade, with two duplication events that resulted in the paralog pairs J2-EJ2 and LIN-RIN in tomato (**Figure S3A-B**). In contrast, the SEP1/2 clade, which comprises SEP1 and SEP2 in Arabidopsis, is reduced to TM29 in tomato, and this contraction to a single SEP1/2 paralog is conserved throughout the *Solanoideae* subfamily of *Solanaceae* (**Figure 5B-C**). Paralog copy number is conserved in the SEP3 clade, which is the ancestral SEP subclade when using the AGAMOUS LIKE6 (AGL6) sister clade as an outgroup, and comprises tomato TM5 and the Arabidopsis ortholog SEP3.

**Figure 5:**
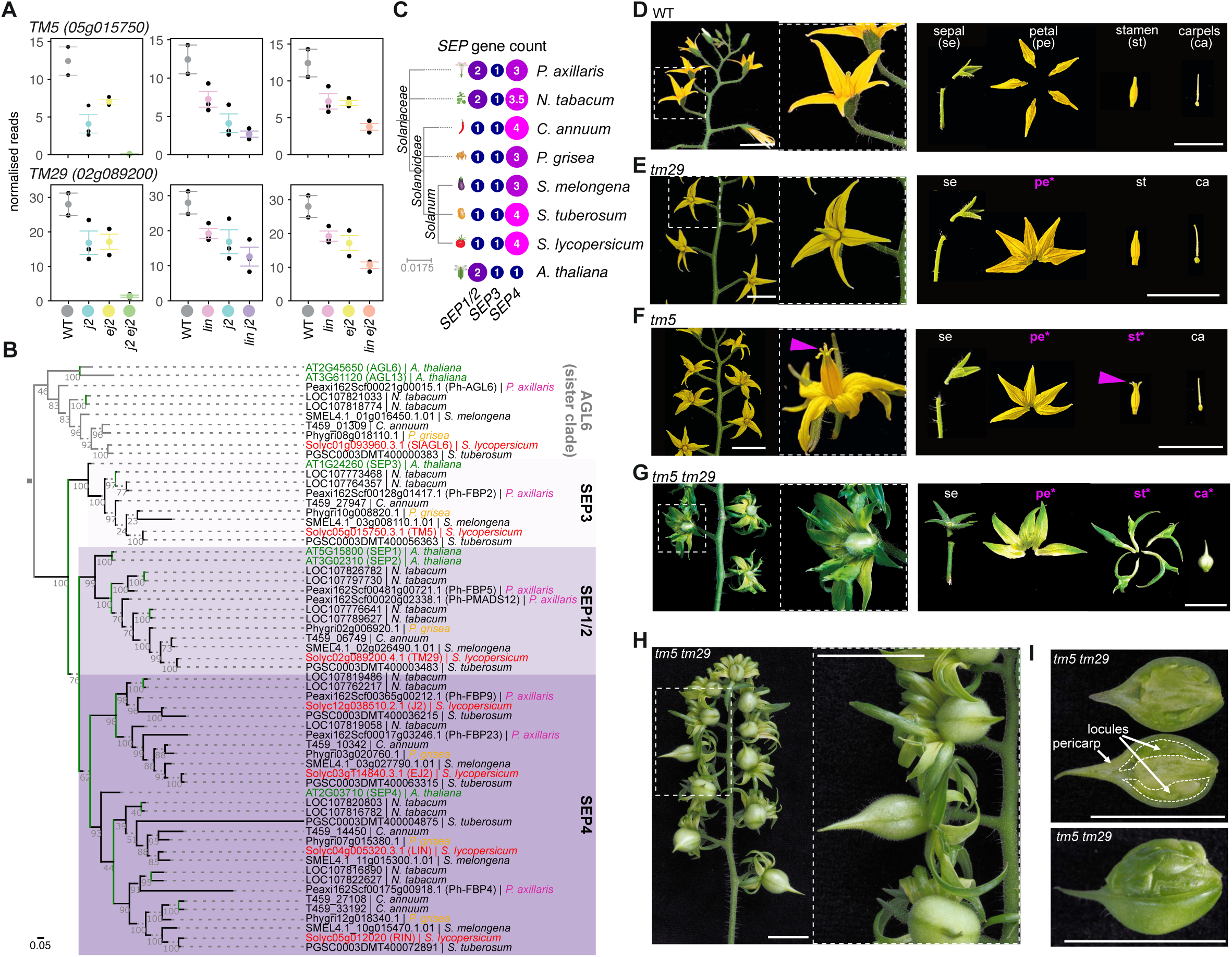
The *SEP1/2* and *SEP3* orthologs *TM29* and *TM5* function redundantly to regulate floral organ identity in tomato. **(A)** Normalized gene expression of *TM5* (top) and *TM29* (bottom) in floral meristem samples from single and double mutant combinations for *j2 ej2*, *lin j2*, and *lin ej2*. Data are represented as mean ± SE. **(B)** Maximum-likelihood tree of SEP proteins in *Arabidopsis* and *Solanaceae*. The AGL6 sister clade was used as outgroup. SEP1/2, SEP3, SEP4 subclades are highlighted in different shades of purple. Tomato, physalis, petunia and Arabidopsis genes are labelled in red, yellow, magenta, and green font, respectively. Numbers at nodes represent ultrafast bootstrap support values from 1,000 replicates. Branch lengths are proportional to the number of amino acid substitutions, branches in green show duplication events, and the scale bar indicates the average number of substitutions per site. **(C)** *SEP* gene counts across distinct subclades in Arabidopsis and *Solanaceae* species shown in the phylogenetic tree in (B). Note that gene content of tetraploid *N. tabacum* is halved to allow comparisons with the other (diploid) genomes. **(D)** Images of a WT inflorescence with a close-up of a single flower (left) and a dissected flower (right), showing the pedicel with calyx (sepals), petals, stamens, and carpels (right). Note the presence of abscission zones on the pedicel of the flower. **(E-G)** Images of the *tm29* (E), *tm5* (F), and *tm5 tm29* (G) mutants as in (D). Note petal fusions in (E-G), bending anthers in (F), and changes in both petal, stamen, and carpel identity in (G) highlighted in magenta font and asterisks. **(H)** Macroscopic image of a *tm5 tm29* double mutant inflorescence with close-up on flowers that produce pseudofruits. **(I)** Macroscopic images of *tm5 tm29* pseudofruits sectioned along the longitudinal axis (top) and with partially removed pericarp (bottom). The pericarp and locules are labelled. Note that locules of pseudofruits are filled with leaf-like organs. Scale bars in (D-I) indicate 1 cm.

### Redundancy between tomato *SEP1/2* and *SEP3* subclades confers the conserved *SEP* function to specify floral organ identity

Since *TM5* and *TM29* were expressed in floral meristems (**Figure S4A**) and synergistically downregulated in *j2 ej2* floral meristem samples (**Figure 5A**), we reasoned that *TM5* and *TM29* may play roles during inflorescence development. Thus, we generated *tm5* and *tm29* mutants and isolated loss-of-function alleles that disrupt the conserved MADS-box domain (**Figure S4B**). Single *tm5* and *tm29* mutants did not show obvious changes in inflorescence architecture (**Figure S4C**). Moreover, we observed only very subtle defects in floral organ development: petals on *tm5* and *tm29* mutants were partly fused, anthers in *tm5* flowers were bending outwards, and *tm5* fruits produced fewer seeds (**Figure 5D-F** and **Figure S4D-F**). We could not confirm the strong floral organ defects of *tm5* and *tm29* single mutants from previous antisense knock-down studies ^44,45^. However, the mild flower defects observed in *tm5* hinted towards unequal redundancy between *TM5* and *TM29* with a more prominent role for the *SEP3* ortholog *TM5* during floral organs specification. Indeed, organ defects were dramatically enhanced in *tm5 tm29* double mutants, which developed flowers with petals and stamen transformed to leaf-like organs (**Figure 5G** and **Figure S4G**). In addition, *tm5 tm29* flowers gave rise to parthenocarpic (seedless) fruits that contained leaf-like organs instead of seeds (**Figure 5I** and **Figure S4H-I**), similar to petunia *sep123* (*fbp2 fbp5*) mutants ^47^. This transformation of floral organs into leaf-like organs in all floral whorls in *tm5 tm29* is reminiscent of the *sep123* triple mutant in Arabidopsis, after which the *SEPALLATA* gene family was named ^24^. However, in contrast to Arabidopsis, loss of *SEP1/2/3* activity in tomato still permits pericarp development in *tm5 tm29* mutants. We concluded that *TM5* and *TM29* act redundantly to confer the conserved *SEP1/2/3* function in specifying floral organ identity across all four whorls. Unequal redundancy between *TM5/TM29* during floral organ specification indicates that they are an ancient pair of paralogs that diverged due to compensatory drift ^3^.

### *SEP4* paralogs retained a function in specifying floral organ identity

The fact that neither single nor double *tm5 tm29* mutants developed branched inflorescences contradicted the hypothesis that synergistic reduction of *TM5* and *TM29* expression is responsible for inflorescence branching in *j2 ej2* mutants. We reasoned that the effect of *TM5* and *TM29* on meristem activity only becomes visible when *SEP4* activity is reduced, so we combined *tm5* and *tm29* with *j2* and *ej2* mutations. Indeed, loss of *J2* or *EJ2* activity in *tm5 tm29 j2* and *tm5 tm29 ej2* triple mutants enhanced the vegetative identity of sepals, petals, and stamen (**Figure 6A-B**). Yet, the most dramatic changes were again observed in the fourth floral whorl. While *tm5 tm29* mutants developed parthenocarpic fruits with leaf-like organs, the additional loss of one *SEP4* paralog in *tm5 tm29 j2* and *tm5 tm29 ej2* triple mutants caused the development of ectopic inflorescences in the center of each flower (**Figure 6A-B** and **Figure S4J-K**). This process is reiterated in each new flower, indicating a reversion from floral to inflorescence meristem identity (**Figure 6C-E**). These results show that when *SEP4* is partially lost in the absence of *SEP1/2/3* (in *tm5 tm29 j2* or *tm5 tm29 ej2*), floral meristems produce three whorls of vegetative organs. Mutant floral meristems fail to terminate in the fourth whorl and instead revert to inflorescence meristem identity, producing an ectopic inflorescence with flowers that reiterate the process. This shows that tomato *SEP4* (*J2* and *EJ2*) can confer the *SEP1/2/3* function in the fourth whorl in the absence of *TM5* and *TM29*, consistent with partial redundancy of *SEP1/2/3/4* during carpel development.

**Figure 6:**
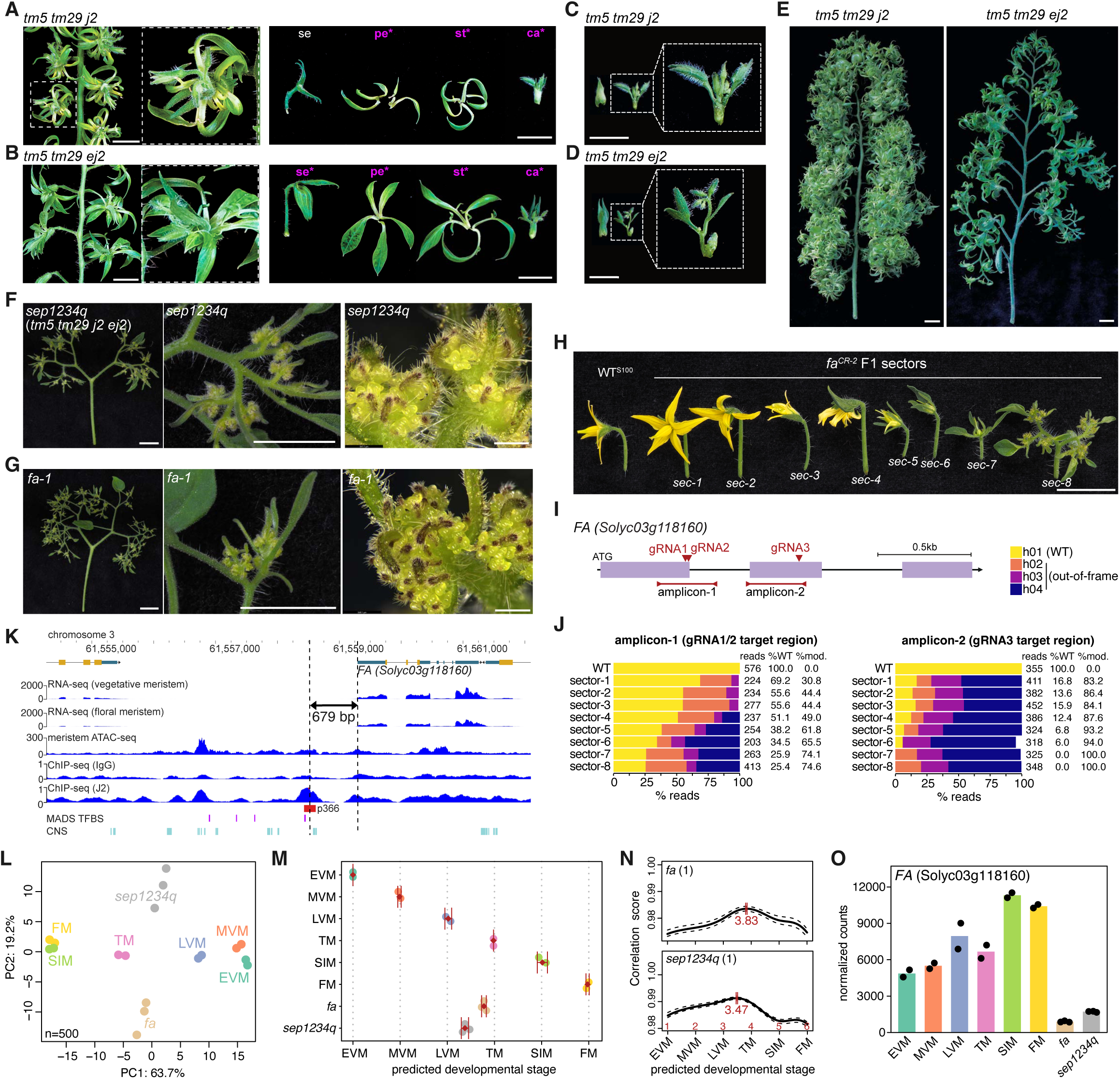
Redundant *SEP1/2/3/4* activity acts on *FALSIFLORA* during the specification of floral organ identities. **(A-B)** Images of the *tm5 tm29 j2* (A) and *tm5 tm29 ej2* (B) triple mutants with a close-up of a single flower (left) and a dissected flower (right), showing the pedicel with calyx (sepals), petals, stamens, and carpels (right). Note changes in both petal, stamen, and carpel identity in (A-B) highlighted in magenta font and asterisks. **(C-D)** Images of dissected of *tm5 tm29 j2* (C) and *tm5 tm29 ej2* (D) triple mutant carpels showing the formation of ectopic inflorescences. **(E)** Images of *tm5 tm29 j2* (left) and *tm5 tm29 ej2* (right) triple mutant inflorescences showing the reiterated formation of ectopic inflorescences in flowers. **(F-G)** Macroscopic (left and middle) and stereomicroscopy (left) images of a *sep1234q* quadruple (F) and *falsiflora* (G) mutant inflorescences with overproliferation of meristem tissue (cauliflower) and bracts. **(H)** Macroscopic images of detached flowers from different sectors (*sec*) of a *fa^CR^* F1 individual. Note the gradual changes in floral organ identify from *sec-2* to *sec-7*, and the cauliflower inflorescence in *sec-8*. **(I)** Schematic representation of *FA* with guide RNAs (gRNA) target positions indicated by arrowheads. Purple boxes represent exons. The Cas9 cleavage sites for gRNAs are indicated with arrowheads. **(J)** Quantification of *FA* haplotype frequencies in different *fa* sectors and a WT control by amplicon deep sequencing. The total number of reads, percentage of wild-type (%WT) and modified (%mod.) reads are shown. Note that haplotype-1 (h01) represents the WT haplotype at both target regions. **(K)** Browser view of the *FA* genomic region. Normalized coverage (CPM) is shown for vegetative and floral meristem RNA-seq, meristem ATAC-Seq, and J2 ChIP-Seq (including IgG control). A significant J2 binding peak (P196) is indicated by a red square. Predicted MADS-box binding sites (MADS TFBS) and conserved noncoding sequences (CNS) are indicted by magenta and cyan squares, respectively. Genomic positions on chromosome 3 (SL4.0) and the distance between the J2 binding site and the transcriptional start site (TSS) are indicated. **(L)** Principal component analyses of the 500 most differential genes in *fa* and *sep1234q* mutant meristems and staged WT meristems (EVM, early vegetative; MVM, middle vegetative; LVM, late vegetative; TM, transition; SIM, sympodial inflorescence; FM, floral). **(M-N)** Developmental stage prediction for *fa* and *sep1234q* mutant meristem tissue from transcriptome data using RAPToR. Age estimates for replicate samples (M) and correlation profile for one *fa* and *sep1234q* replicate each (N) are shown. Diamonds and error bars in (M) indicate mean values and standard deviation. Red bars indicate estimation confidence intervals, and black dotted lines indicate the 95% interval of bootstrap correlation with the reference. The sample age estimate is displayed below the interval. Estimated reference stage is indicated in red font. **(O)** Expression (normalized read counts) of *FA* in staged WT meristems and *fa* and *sep1234q* mutant meristem tissue. Scale bars indicate 1 cm (A-E, F-G (left), H), 0.5 cm (F-G middle), 500 µm (F-G right).

### *SEP1/2/3* paralogs confer a minor function in driving meristem transitions to floral identity

We further reduced *SEP4* activity in the absence of *SEP1/2* or *SEP3* paralogs by isolating *tm29 j2 ej2* (*sep1/2 sep4^partial^*) and *tm5 j2 ej2* (*sep3 sep4^partial^*) triple mutants. The triple mutants developed inflorescence branching patterns reminiscent of *j2 ej2* (**Figure 2B** and **Figure S5A**). In addition, loss of *TM5* (*SEP3*) or *TM29* (*SEP1/2*) increased the occurrence of vegetative organs (bracts) in the inflorescence and defects in floral organ identity (**Figure S5A, B**). As a result, *tm5 j2 ej2* mutants developed petals instead of stamens, indicating that loss of *J2* and *EJ2* further enhances effects in *tm5* on stamen identity. We also isolated the *tm5 tm29 j2 ej2* quadruple (*sep1234q*) mutant, which developed highly branched inflorescences that failed to differentiate flowers and instead developed modified leaves (bracts) and overproliferative meristematic tissue (**Figure 6F**). This severe loss of floral identity from the combined mutation of *TM29* (*SEP1/2*), *TM5* (*SEP3*), and *J2/EJ2* (*SEP4*) shows that *SEP1/2/3* paralogs confer floral meristem identity when *SEP4* dosage is reduced. In conclusion, *SEP1/2/3* paralogs retained a redundant function with *SEP4* paralogs in promoting meristem maturation towards floral identity.

### *SEP1/2/3/4* redundancy acts on the conserved floral identity gene *FALSIFLORA*

The inflorescences of *sep1234q* mutants closely resembled those of the historical mutant *falsiflora (fa)* ^15^, which lacks the activity of the tomato ortholog of Arabidopsis *LFY*, a floral identity gene that is deeply conserved across flowering plants ^14,16,48^. Hence, we generated *fa* mutants in cherry tomato (cv. Sweet-100) using CRISPR–Cas and observed the expected *fa* phenotypes in multiple first-generation (T0) plants (**Figure 6G** and **Figure S5D-F**). Due to high Cas9 editing activity, we recovered only one out of eleven *fa^CR^* T0 individuals that developed a few fertile flowers with pollen for backcrossing to the WT. Cas9-free *fa-null/+* F1 heterozygotes did not develop obvious floral organ or inflorescence defects (**Figure S5G**), confirming the recessive nature of the *fa* mutant ^15^. However, we identified an F1 individual that retained Cas9 activity and developed distinct sectors with abnormal phenotypes ranging from floral organ defects to *fa-null* inflorescences that overproliferated bracts and meristematic tissue (**Figure 6H** and **Figure S5H**). The distinct phenotypic sectors suggested that Cas9 had induced additional *fa* mutations, leading to gradual reductions in *FA* dosage and a spectrum of floral organ and inflorescence defects. We quantified the frequency of *FA* mutations in individual flowers from sector 1(sec-1) to sec-7 or inflorescence tissue (sec-8) by amplicon deep sequencing (**Figure 6I**). Remarkably, the frequency of the functional *FA* (WT) haplotype indeed decreased with phenotypic severity, from sectors with mild floral organ defects to sectors with strong organ defects and branched inflorescences, which contained only out-of-frame *fa* mutations (**Figure 6J**, **Figure S5I**, and **Table S11**). These results show that gradual reductions in *FA* gene dosage can lead to both floral organ defects and inflorescence branching. Given the phenotypic similarities between *fa* and *sep1234q* mutants, we reasoned that *SEP1/2/3/4* activity is required for maintaining *FA* expression in reproductive meristems, which guides meristem maturation towards floral fate. This model is supported by a J2 binding site upstream of *FA* with two CarG-box elements (**Figure 6K**) ^41^ and a recent report that deletion of this site leads to floral organ defects and mild inflorescence branching ^49^.

To investigate the functional relationship between *SEP1/2/3/4* and *FA*, we profiled gene expression in *fa* and *sep1234q* inflorescences by RNA-seq and compared mutant transcriptomes with staged WT meristem profiles ^5^. Principal component analyses of the 500 most variable genes resolved subsequent stages of meristem maturation on the first component (PC1) (**Figure 6L**). To estimate the developmental stage of *fa* and *sep1234q* meristem tissue, we applied an established framework ^50^ and staged mutant transcriptomes on the WT meristem maturation trajectory as reference (**Figure S6** and **Methods**). This analysis predicted *fa* and *sep1234q* near late vegetative (LVM) and transition (TM) meristem stages, suggesting that *fa* and *sep1234q* meristems revert to a maturation stage that precedes the transition to reproductive development (**Figure 6M-N**). In addition, *FA* transcript levels were strongly reduced in *sep1234q* meristems, consistent with a model in which *SEP1/2/3/4* activity is required to maintain *FA* expression, which is essential to acquire reproductive meristem identity (**Figure 6O**). Together, these results indicate that incomplete functional divergence of *SEP* genes ensures redundancy on the dose sensitive *AN-FA* module, which is conserved across flowering plants to drive meristem maturation during inflorescence and flower development ^48^.

## DISCUSSION

### Synergy arises from redundancy on conserved developmental regulators

In this study, we investigated the molecular principles of genetic synergy by dissecting genetic interactions between closely related *SEP4*-clade MADS-box genes in the model crop tomato. Previous studies showed synergy between the *SEP4*-clade paralogs *J2*, *EJ2*, and *LIN* on the suppression inflorescence branching ^12,51^. Here, we found that *SEP4* synergy on gene expression programs underlying inflorescence branching is more pronounced between the direct paralogs *J2* and *EJ2* than with the distal paralog *LIN*, indicating that synergistic effects decline with evolutionary distance. Transcriptome analyses show that *SEP4* synergy acts through a defined set of genes in floral meristems including *AN*, a conserved floral identity gene and ortholog of Arabidopsis *UFO* ^17,48^. By integrating data from previous studies ^41,42^, we provide evidence that SEP4 paralogs directly activate *AN* expression by binding an enhancer element ∼3.5 kbp upstream of *AN* when meristems transition to floral identity, a critical window in which inflorescence architecture is predetermined ^5,12,13^. Further, we found that synergy between four *SEP1/2/3/4* paralogs acts through the expression of *FA*, a second deeply conserved floral identity gene and ortholog of Arabidopsis *LFY* ^14,20,48^. Remarkably, loss of *SEP1/2/3/4* activity leads to lower *FA* transcript levels, and lowering *FA* gene dosage recapitulates the phenotypic spectrum arising from gradual reduction of *SEP1/2/3/4* dosage. These results indicate that *SEP1/2/3/4* synergy converges to maintain *FA* dosage. We found evidence that SEP1/2/3/4 factors directly regulate *FA* by binding to an upstream *cis*-regulatory element, a model supported by published ChIP-Seq data ^41^ and by CRISPR-Cas deletion of this element, which leads to inflorescences branching and homeotic floral organ conversions ^49^. Thus, residual *SEP1/2/3/4* redundancy acts on the *FA-AN* module to drive meristem transitions to reproductive development, a function that remained conserved when *SEP* genes functionally diverged. Together, these findings show that gene synergy emerges as a relic of functional redundancy.

### Feedback regulation between paralogs calibrates their gene dosage

We speculate that transcriptional feedback regulation between *SEP* paralogs coordinates subsequent stages of meristem maturation by maintaining a critical dosage of the *FA-AN* module. *SEP4* synergy influences the expression of two closely related *SEP1/2/3* genes, leading to a dose-dependent downregulation of *SEP1/2/3* paralogs in floral meristems when *SEP4* activity is reduced. Whether SEP4 proteins directly activate *SEP1/2/3* paralogs remains to be determined since we could not identify J2 binding sites near *TM5* and *TM29*, suggesting either an indirect relationship or a direct control over longer genomic distances. Nevertheless, this transcriptional feedback between *SEP4* and *SEP1/2/3* paralogs appears essential for calibrating *SEP* dosage to maintain a critical expression level of the *FA*-*AN* module in meristems during the acquisition of floral identity. The Arabidopsis orthologs of the transcription factor FA (LFY) and the F-box protein AN (UFO) form a heteromeric transcriptional complex that regulates targets which LFY homomers cannot bind ^39^. In tomato, *FA* is expressed throughout meristem maturation including developmental stages that precede the floral transition. In contrast, *AN* is specifically expressed during the floral stage of meristem maturation. Thus, FA homomers likely confer the *FA* function in promoting the transition to flowering, consistent with a flowering delay observed in *fa* but not in *an* mutants ^14,20^. The activation of *SEP4* paralogs at the floral transition precedes *AN* expression in floral meristems, which is diminished when *SEP4* activity is reduced. *SEP4* function is thus required for *AN* expression and the formation of putative FA-AN heterocomplexes that drive meristem maturation post transition towards floral identity. *SEP1/2/3* paralog expression is instead activated in floral meristems during the initiation of floral organs ^9^, and a critical *SEP* dosage is essential during these maturation stages to maintain *FA* expression. Progressive loss of *SEP* activity in *tm5 tm29* (*sep123*) and *tm5 tm29 j2 ej2* (*sep1234q*) mutants causes homeotic organ conversions and loss of floral meristem determinacy. Similarly, quantitative reductions in *FA* dosage cause floral organ conversions and loss of floral meristem identity ^49^, and *FA* expression is substantially reduced in *sep1234q* mutant meristems. Thus, transcriptional feedback regulation between *SEP1/2/3/4* paralogs ensures a critical *SEP* dosage to maintain *FA* and *AN* expression after the floral transition that is required for accurate floral meristem patterning and floral organ differentiation.

### Gene dosage effects maintain redundancy but constrain functional divergence

Multiple duplication events led to the expansion of the *SEP* gene family and allowed *SEP* paralogs to functionally diverge and acquire new functions ^23,32^. We show that *SEP* redundancy is conserved during the transition of meristems to floral identity, seen in reversion of floral meristems to vegetative identity in *sep1234q* mutants. Redundant *SEP* roles during floral meristem development have been described in other species ^32,33^, which indicates that the *SEP* function in guiding meristem maturation towards floral identity is conserved. *SEP3*, which is the ancestral paralog and conserved to one copy per diploid genome in most dicot species, thus evolved the derived and more specialized function in floral organ identity. Nevertheless, the dosage sensitivity of *SEP* genes during inflorescence and flower development imposed a constraint to the functional divergence of the *SEP* family. The molecular basis that underlies incomplete divergence of *SEP* paralogs remains to be determined. Divergent expression patterns from *cis*-regulatory changes may led to functional divergence without unbalancing *SEP* dosage, and expression atlas data for *SEP* genes ^5,9^ hints towards regulatory sequence evolution although more detailed analyses of cell type-specific expression patterns are required. Future analyses of non-coding sequences may reveal *cis*-regulatory regions and transcriptional enhancers that control spatiotemporal *SEP* expression patterns underlying both divergent and redundant functions. In summary, our work demonstrates that dosage sensitivity of paralogous genes can impose an evolutionary constraint to the capacity of gene families to fully diverge in function.

## METHODS

### Plant material and growth conditions

Wild-type seed for the *S. lycopersicum* cv. M82 (LA3475) and *S. lycopersicum* cv. Sweet-100 double-determinate ^40^ were from our own stocks. Mutant seed for *j2*, *ej2* and *lin* single and combinatorial mutant combinations in the M82 background were from our own stocks ^12^. For phenotypic analyses, seeds were germinated on soil in 96-well plastic trays and grown under long-day conditions (16-h light/8-h dark) in a greenhouse supplemented with artificial light from LED panels (∼250 μmol m^−2^ s^−1^), oscillating day-night temperature (25 °C day /20°C night), and relative humidity of 50%–60%. Plants were grown in 5 L pots (2 per pot) under standard fertilizer regime and drip irrigation. For meristem analyses, seeds were pre-germinated on moist filter paper at 28°C in the dark for 72 hours. Germinated seedlings with similar radicle length were transferred to soil in 96-well plastic flats and grown in the greenhouse on flooding tables.

### Plant phenotyping

To quantify flowering time, the number of leaves before the emergence of a floral meristem was counted. A minimum of 9 plants was included for each of the genotypes. Additionally, the number of seedlings in vegetative, transition and floral meristem stage was assessed for each of the genotypes 20 days after sowing in a minimum of 22 plants per genotype. To quantify number of flowers per inflorescence, flowers and floral buds were counted on five inflorescences per plant (excluding the first inflorescences) and 8 replicate plants per genotype. Inflorescence branching was quantified by counting the number of branch points in five inflorescences per plant and 8 replicate plants per genotype. Whenever that number exceeded 22 branching points the inflorescence was considered branched but “uncountable”. All phenotyping experiments were performed for all the genotypes in parallel on a single day.

### Stereoscopy and photography

Seedlings grown in 96-well trays were harvested individually and all primordia except for the last two were removed by hand or with a needle. Z-stack images of developing meristems were acquired in different maturation stages spanning from vegetative to inflorescence using a Leica M205 FCA stereomicroscope with a Leica 10450028 objective connected to a Leica DFC7000 T lamp (Leica Microsystems, Wetzlar, Germany). Photographs of inflorescences were taken using either an Olympus OM-D camera with an ED 14-42 mm f/3.5-5.6 EZ objective or an iPhone11 equipped with a 12 MP wide-angle camera.

### CRISPR-Cas

CRISPR–Cas9 mutagenesis in tomato was performed as previously described ^52–54^. Briefly, guide RNAs (gRNAs) were designed using CRISPOR ^55^ and the Sweet-100v.2.0 reference genome ^40^. Binary vectors for CRISPR-Cas mutagenesis were assembled using the Golden Gate cloning system as previously described ^52,56,57^. Briefly, gRNA target sequences (gRNAs) were amplified using KOD Hot Start DNA Polymerase (Merck Millipore) using target-specific forward primer, a universal reverse primer and the pICH86966::AtU6p::sgRNA vector (Addgene 46966) as template. The gRNAs were combined with the AtU6p promoter in Level1 vectors, and assembled with Nos::NptII::ocs (addgene #51144) and SlUbi::SpCas9-P2A-GFP::nos ^52^ in Level2 binary vectors. Plasmids were verified by Sanger sequencing and transformed into *Agrobacterium tumefaciens* (AGL-1) by electroporation using a Bio-Rad GenePulser II (1 mm cuvettes, 1.8 kV, 25 µF, 200 Ω, ∼5 ms pulse). Constructs were transformed into the double-determinate cherry tomato cultivar Sweet-100 or M82 by *Agrobacterium tumefaciens*-mediated transformation as previously described. CRISPR–Cas9 editing was verified in the T0 generation by genotyping, Sanger sequencing or amplicon deep sequencing as described ^52^, and transgene-free mutant plants were isolated in the T1. *J2*, *EJ2* and *LIN* were targeted simultaneously with a multiplex construct containing two gRNAs per gene. *TM5* and *TM29* were targeted with two independent constructs with two gRNA per gene, and mutations were combined by crossing T1 or T2 individuals. *FA* was targeted with three gRNA sequences. All gRNAs used in this study are listed in **Table S12**.

### Genotyping

Frozen leaf or cotyledon tissue was homogenized with two metal beads at a frequency of 18–20 Hz for 1 min in a mix mill. Genomic DNA was extracted using extraction buffer (0.1 M Tris-HCl pH 9.5, 0.25 M KCl, 0.01 M EDTA) by vigorous vortexing. Cell extracts were transferred to new multi-well plate, incubated at 95°C for 10 minutes, and subsequently cooled at 4°C for 5 min. Equal volume of 3% (w/v) BSA was added to the lysate, mixed vigorously by vortexing, and centrifuged at 3,700 rpm for 15 minutes. Target regions were amplified with gene-specific primers and products were separated on agarose gels. For CAPS analyses, products were digested using restriction enzymes (NEB). For Sanger sequencing, PCR products were purified with ExoSap (Thermo). All primer sequences and genotyping information are listed in **Table S13**.

### Amplicon deep sequencing

Genetic sector analyses were performed using amplicon deep sequencing as previously described ^52,58^. Briefly, gDNA was extracted from individual flowers and gRNA target regions were amplified using gene-specific primers (**Table S13**) with common adapter sequences. Gene-specific amplicons were diluted ten-fold in H_2_O and used as template for the addition of sample-specific indexes using adapter primers with unique barcodes as described before ^40,58^. Equal volumes of each indexed amplicon reaction were pooled and subsequently purified using SPRIselect beads (Beckman Coulter). A single Illumina sequencing library was prepared using the xGen DNA Library Prep MC Kit (Integrated DNA Technologies) and sequenced on the MiSeq System (Illumina) at the Genome Technologies Facility (GTF) at UNIL. Raw reads were analyzed using the S100 reference genome^40,52^ and a previously published pipeline ^52^ based on Trimmomatic (v0.39) ^59^, FASTX-Toolkit (v0.0.14) ^60^, PEAR (v0.9.6), HISAT2 (v2.2.1) ^61^, SAMtools (v.1.17) ^62^, SMAP ^63,64^, and MAFFT (v7.505) ^65^, retaining haplotypes with a minimum frequency of 5%. Haplotypes were categorized in “WT” (no insertions or deletions), “in-frame” (total InDel size a multiple of three or zero), “out-of-frame” (total InDel size not a multiple of three nor zero). Haplotype frequencies and sequences were visualized in R ^66,67^ and Benchling ^68^, respectively.

### Cis-regulatory sequence analyses

Accessible chromatin regions (ACRs) were identified using published ATAC-seq data ^69^. Binding site for J2 were identified using published J2 ChIP-seq data ^41^. Conserved non-coding sequences (CNS) were retrieved from the conservatory database ^70^. MADS-box transcription factor binding sites were predicted on the SL4.0 reference genome ^71^ by FIMO/MEME Suite ^72^ using default parameters and the non-redundant set of profiles for MADS box factors from the JASPAR CORE database (release 2020) ^43^(**Table S14**). Data was visualized in jbrowse2 ^73^.

### Meristem dissection, RNA extraction, library preparation and sequencing

Thirteen days after sowing, three representative seedlings per 96-well flat were dissected to determine the average developmental stage of the seedling population. When representative seedlings had reached the transition or floral meristem stage of meristem maturation ^5^, seedings with similar number of leaves were rapidly harvested starting at 13:00 hrs for a maximum of 2 hrs to minimize circadian effects. After removal of leaves, shoot apices were fixed in ice-cold acetone and vacuum infiltrated for 45 min at 4 °C. Meristems were micro-dissected using a sharp syringe needle under a Leica MS5 stereomicroscope connected to a Leica CLS 100X lamp (Leica Microsystems, Wetzlar, Germany). Dissected meristems of the same genotype and morphological stage were pooled in 2 ml round-bottom RNAse-free Eppendorf tubes (Eppendorf, Hamburg, Germany) filled with ice-cold acetone, and stored at -70 °C. RNA was extracted from pools of 18-38 meristems and three replicate pools per genotype-stage combination using the Arcturus™ PicoPure™ RNA isolation kit (Applied Biosystems) according to the manufacturer protocol with minor modifications. Volumes of extraction buffer and 70% EtOH were increased 2.2-fold and RNA was treated with DNAse to remove contaminating gDNA using the RNAse-Free DNase kit (Qiagen). Sequencing libraries were prepared from 300 ng total RNA as starting material at the Genomic Technologies Facility (GTF) of the University of Lausanne. Indexed libraries were prepared using the TruSeq Stranded mRNA Library Prep kit from Illumina according to the manufacturer’s instructions. Fragment size and concentration were assessed with a Bioanalyzer. Libraries were sequenced on two Illumina NovaSeq6000 lanes to produce between 15,819,728 and 22,075,610 SR100 reads per sample (**Table S15**).

### Read quality control and alignment

Read quality of raw reads was assessed using the FastQC (v0.11.7) ^74^. Illumina TruSeq adaptors were trimmed with Trimmomatic (v0.36; with parameters SE -phred33 ILLUMINACLIP:TruSeq3-SE.fa:2:30:10 LEADING:3 TRAILING:3 SLIDINGWINDOW:4:15 MINLEN:36) ^59^. Trimmed reads were aligned to the SL4.0 reference genome^71^ using STAR (v2.7.8a; with parameters --runMode alignReads -- outFilterType BySJout -- outFilterMultimapNmax 20 -- outMultimapperOrder Random --alignSJoverhangMin 8 --alignSJDBoverhangMin 1 --alignIntronMin 20 –alignIntronMax 1000000 --alignMatesGapMax 1000000 --outSAMtype BAM Unsorted) ^75^. Alignments were sorted and indexed using SAMtools (v.1.17) ^62^ and gene expression was quantified as unique reads aligned to the SL4.0 gene annotation (ITAG4.0) ^71^ using HTSeq-count (v.0.11.2; with parameters --format=bam --order=pos --stranded=no --type=exon --idattr=Parent) ^76^.

### Differential expression (DE) analyses and identification of non-additive expression pattern

Transcriptomic data were processed using the edgeR (v4.4.2) ^77^ and limma (v3.62.2) ^78^ packages for differential expression analysis. Synergistic expression patterns were identified following the analysis framework described by Schrode et al. ^37^. In brief, lowly expressed genes were filtered retaining only those with ≥ 10 reads in at least 2 samples. Counts were normalized using the trimmed means of M-values (TMM) method and transformed into log2 counts per million (logCPM) with the voom function() which also estimates mean-variance relationships and assigns precision weights for downstream linear modelling ^37,78^. For each genotype (e.g. WT, *j2, ej2, j2ej2)*, a design matrix was constructed, and gene-wise linear models were fitted using the lmFit() function. For each set of singles and corresponding double mutant contrasts were defined as follows:

- Additive expectation=(*j2*−WT)+(*ej2*−WT)
- Combinatorial effect=(*j2ej2*−WT)
- Synergy (interaction)=(*j2ej2*−WT)−[(*j2*−WT)+(*ej2*−WT)]

For each gene moderated t-statistics were calculated using the empirical Bayes procedure implemented in eBayes(), shrinking gene-wise variance estimates toward a common prior, stabilizing the standard errors (SE). Statistical significance was defined as genes with 95% confidence intervals excluding zero after Benjamini–Hochberg (BH) multiple-testing correction (FDR < 0.05). Each logFC estimate was tested as LogFC±1.96×SE. Based on the calculated significance, genes were then classified as:

- Non-additive: when the synergy contrast was significant (FDR < 0.05). It can be:

- Synergistic: when the double mutant effect exceeded the additive expectation in the same direction (i.e. more up or more down than predicted).
- Less-than-additive: when the double mutant effect was weaker than expected.
- Additive when the double mutant was significantly different from control (FDR < 0.05) but the synergy contrast was not significant.

To further evaluate the robustness of synergy detection, the proportion of non-null hypotheses (π₁) across all synergy tests was estimated using the Storey–Tibshirani qvalue method ^79^. Median standard error (SE) of contrasts was used to estimate the probability of detecting deviations from additivity at varying effect sizes and sample numbers^37^. Differentially expressed genes (DEGs) per genotype (single vs. control or double vs. control) were determined with a threshold of |log2FC| ≥ 0.58 (∼1.5-fold change) and FDR < 0.01.

Multidimensional scaling (MDS) analyses were conducted using CPM values on: all expressed genes, 2583 genes dynamically expressed during meristem maturation ^13^ and in 227 transition marker and 241 floral meristem marker genes, in the transition and floral stage datasets, respectively ^12^. The samples linej2_TM_3 and M82_FM_1 were excluded from the differential gene expression analyses due to outlier behavior in the transition marker and floral marker MDS analyses (**Figure S2**). Gene normalized z-scores were calculated with the formula z = (x -μ) / σ, where x is the CPM value for a specific gene, μ is the mean of CPM values across samples, and σ is the standard deviation. All data visualization and statistical analyses were conducted in R ^66,67^. Gene normalized z-scores were plotted as heatmaps using pheatmap (v.1.10.12) ^80^ and dotplots using ggplot2 package (v3.5.1) ^81^. Counts per million (CPM) were plotted in dotplots using ggplot2 (v3.5.1) ^81^. Overlaps of DEGs were visualized using the eulerr package (v7.0.2) ^82^.

### RNA-seq analysis *sep1234q* and *fa* inflorescence tissue

Meristem tissue for mRNA extraction was collected at 13:00 hrs from mature *sep1234q* and *fa* plants grown in a greenhouse. Per genotype, 3 individual plants served as biological replicates, 3 inflorescences (∼5 mm) were pooled per replicate, and flash-frozen in liquid nitrogen. Total RNA was extracted using the RNeasy Plant Mini Kit (QIAGEN) following the manufacturer’s protocol and assessed for quality using a Bioanalyzer. Sequencing libraries were prepared from 100 ng of total RNA at the Genomic Technologies Facility (GTF), University of Lausanne. In brief, indexed libraries were generated using the WATCHMAKER mRNA Library Prep Kit (Watchmaker Genomics) according to the manufacturer’s instructions, and library fragment size distribution and concentration were evaluated using a Bioanalyzer. Libraries were sequenced on two Element AVITI lanes, yielding between 17,145,097 and 20,335,750 paired-end 150 bp reads per sample.

Read quality controls, alignments and counts were produced as described in the previous section. Lowly expressed genes were discarded by retaining only genes with ≥10 counts in at least 3 samples. Differential expression analysis and variance-stabilizing transformation (VST) was conducted in DESeq2 (v1.46.0)^83^. The transformed expression matrix was corrected for batch effects (WT samples ^5^ versus mutant (*sep1234q* and *fa*) samples) using the removeBatchEffect function from the limma (v3.62.2) ^78^. Estimation of developmental stage was conducted using real-age prediction from transcriptome staging on reference (RAPToR, v1.2.0) ^50^. A reference model was built on the staged WT meristem data using the functions ge_im() (formula = X ∼ s(time, bs = “ts”, k = 5), nc = 3) and make_ref() (n.inter = 100). The WT reference model was used to stage mutant samples using the ae() function. Correlation profiles and stage estimates were visualized with plot() and plot_cor(), respectively, and mean stage estimates in ggplot2 (v3.5.2).

### Gene ontology enrichment analysis

Gene ontology enrichment analyses for biological processes were performed among differentially expressed genes using a functional annotation database for the ITAG4.0 tomato genome from PLAZA ^84^ and the ClusterProfiler package ^85^ with a pvalueCutoff = 0.05 and pAdjustMethod = BH. All annotated genes in the tomato genome (ITAG4.0) were used as background.

### Phylogenetic analysis

To reconstruct the phylogenetic tree of the SEP family across angiosperm species, a total of 22 proteomes were collected from the Orthologous MAtrix (OMA; release 12/2024) ^46^ database and the Conservatory CNS project ^70^ (**Table S10**). Splice variants were removed to only retain primary protein isoforms and reduce redundancy. Hierarchical Orthologous Groups (HOGs, gene families) were obtained using FastOMA (v1.2.0) ^86^ and a species tree from the NCBI Taxonomy Database ^87^ as the guide phylogeny (**Supplementary Data 1**). The SEP and AGL6 gene families were identified using the proteins from *S. lycopersicum* and *A. thaliana* (**Table S9**). The 135 protein sequences of this gene family were aligned with MAFFT (v7.505) ^65^, MUSCLE (v3.8.1551) ^88^ and Kalign (v3.3.2) ^89^ using default parameters. Resulting alignments were combined to a consistency-based alignment using T-Coffee (v13.46.0.919e8c6b) ^90^ ^91^. The maximum likelihood (ML) tree was inferred using IQ-TREE (v1.6.12) with 1000 ultrafast bootstrap replicates ^92^. This tree was pruned using the ETE4 (Environment for Tree Exploration) ^93^ toolkit to include only representatives of the Solanaceae family and Arabidopsis. To identify gene duplications, the gene trees were reconciled with the species tree via the species-overlap method ^94^ using the ETE4 toolkit.

## DATA AVAILABILITY

Raw Illumina sequence data generated in this study will be made available on SRA under the BioProject PRJNA1348489 upon publication. Seeds are available on request from S. Soyk.

## ACKNOWLEDGEMENTS

We thank all members of the Soyk lab for helpful discussions and comments on the manuscript; J. Marquis and J. Weber from the Genomic Technologies Facility (GTF) of the UNIL for support with sequencing; B. Tissot, L. Nerny, L. Keel, T. Stupp, T. Bovey, E. Inacio Martins, and L. Héau for support with plant care; L. Lebeigle for technical support; R. Dreos for bioinformatics support. This work was supported by the University of Lausanne, the European Research Council (ERC) under the European Union’s Horizon 2020 research and innovation programme (ERC Starting Grant “EPICROP” Grant No. 802008) to S.So., the Swiss National Science Foundation (SNSF) under an Eccellenza Professorial Fellowship (Grant No. PCEFP3_181238) to S.So., and an UNIL Interdisciplinary Project Grant to N.Gl. and S.So.

## AUTHOR CONTRIBUTIONS

N.G. and S.So. conceived the project. N.G., Y.X, S.So. designed and planned experiments. N.G., Y.X, E.L. and S.So. performed experiments and collected data. N.G., Y.X, E.L., I.J. and S.So. analyzed data. N.G. and S.So. acquired project funding. N.G. and S.So. wrote the first draft of the manuscript. All authors read, edited, and approved the manuscript.

---

**Figure S1:**
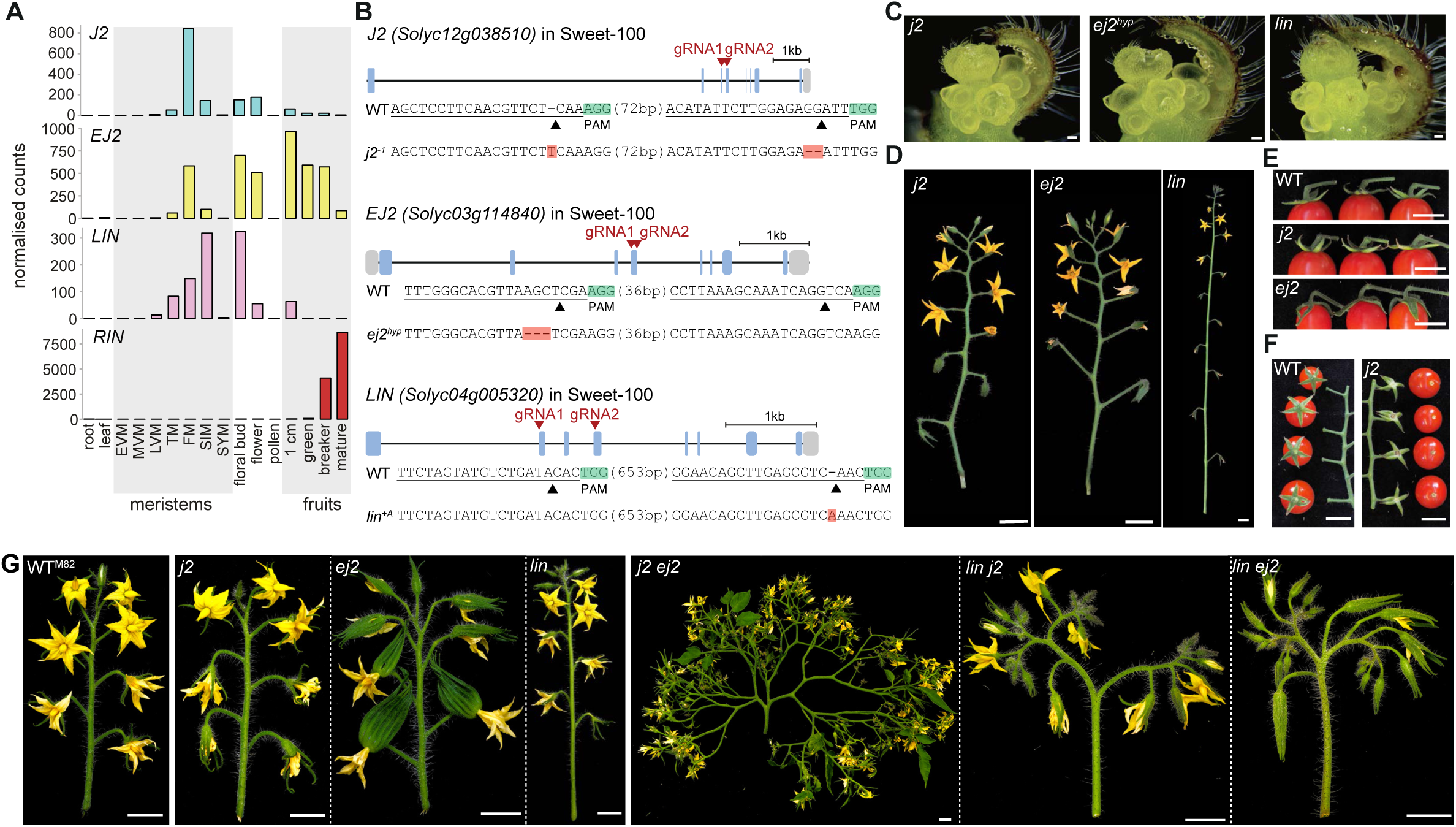
The SEP4 paralogs J2, EJ2, and LIN have unequal contribution to inflorescence development. **(A)** Normalized read counts (transcripts per million, TPM) for the tomato SEP4 genes J2, EJ2, LIN, and RIN in different tissues and meristem stages (EVM, early vegetative; MVM, middle vegetative; LVM, late vegetative; TM, transition; FM, floral; SIM, sympodial inflorescence; SYM, sympodial shoot meristem). **(B)** Schematic representation of J2 (top), EJ2 (middle), and LIN (bottom) with guide RNAs (gRNA) target positions indicated by arrowheads. Blue boxes represent exons and grey boxes represent UTRs (untranslated regions). The Cas9 cleavage sites for gRNAs are indicated with arrowheads. Mutant allele sequences identified by Sanger sequencing are shown with mutated nucleotide positions highlighted with red background. The Cas9 cleavage sites for guide RNAs are indicated with arrowheads and PAMs are marked in green. PAM, protospacer adjacent motif. **(C)** Stereomicroscopy images of a j2 (left), ej2 (middle), and lin (right) single mutant shoot apices with developing primary inflorescences. **(D)** Images of detached inflorescences of the j2 (left), ej2 (middle), and lin (right) single mutant. **(E)** Images of detached fruits with calyx of the WT (top), j2 (middle) and ej2 (bottom). Note the elongated sepals in ej2. **(F)** Images of inflorescences and hand-picked fruits of the WT and j2. Note the lack of fruit abscission at the pedicel in j2. **(G)** Detached inflorescences from the M82 wild-type (WT^M82^) and the mutant collection for j2, ej2, and lin single and double mutants in the tomato cultivar M82. Scale bars indicate 100 µm (C) and 1 cm (D-G).

**Figure S2:**
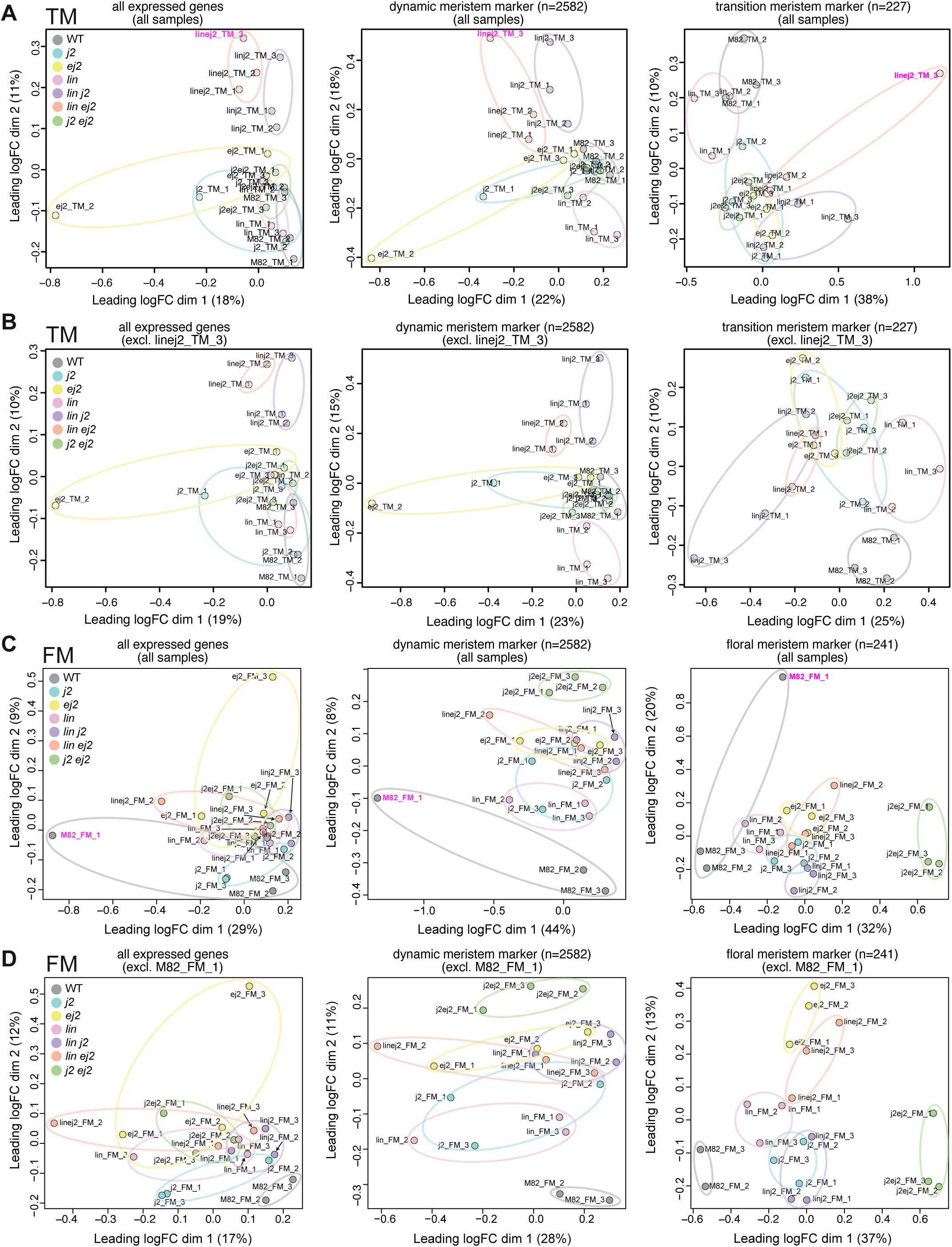
Multidimensional scaling (MDS) plots for gene expression profiles in transition and floral meristem samples. **(A)** MDS plots based on normalized counts for all expressed genes (left), genes dynamically expressed across stages of meristem maturation (middle), and transition stage marker genes (right) in all transition meristem (TM) samples. The outlier sample linej2_TM_3 is highlighted in magenta. **(B)** MDS plots as in (A) for transition meristem (TM) samples excluding linej2_TM_3 outlier sample. **(C)** MDS plots based on normalized counts for all expressed genes (left), genes dynamically expressed across stages of meristem maturation (middle), and floral stage marker genes (right) in all floral meristem (FM) samples. The outlier sample M82_FM_1 is highlighted in magenta. **(D)** MDS plots as in (C) for floral meristem (FM) samples excluding M82_FM_1 outlier sample.

**Figure S3:**
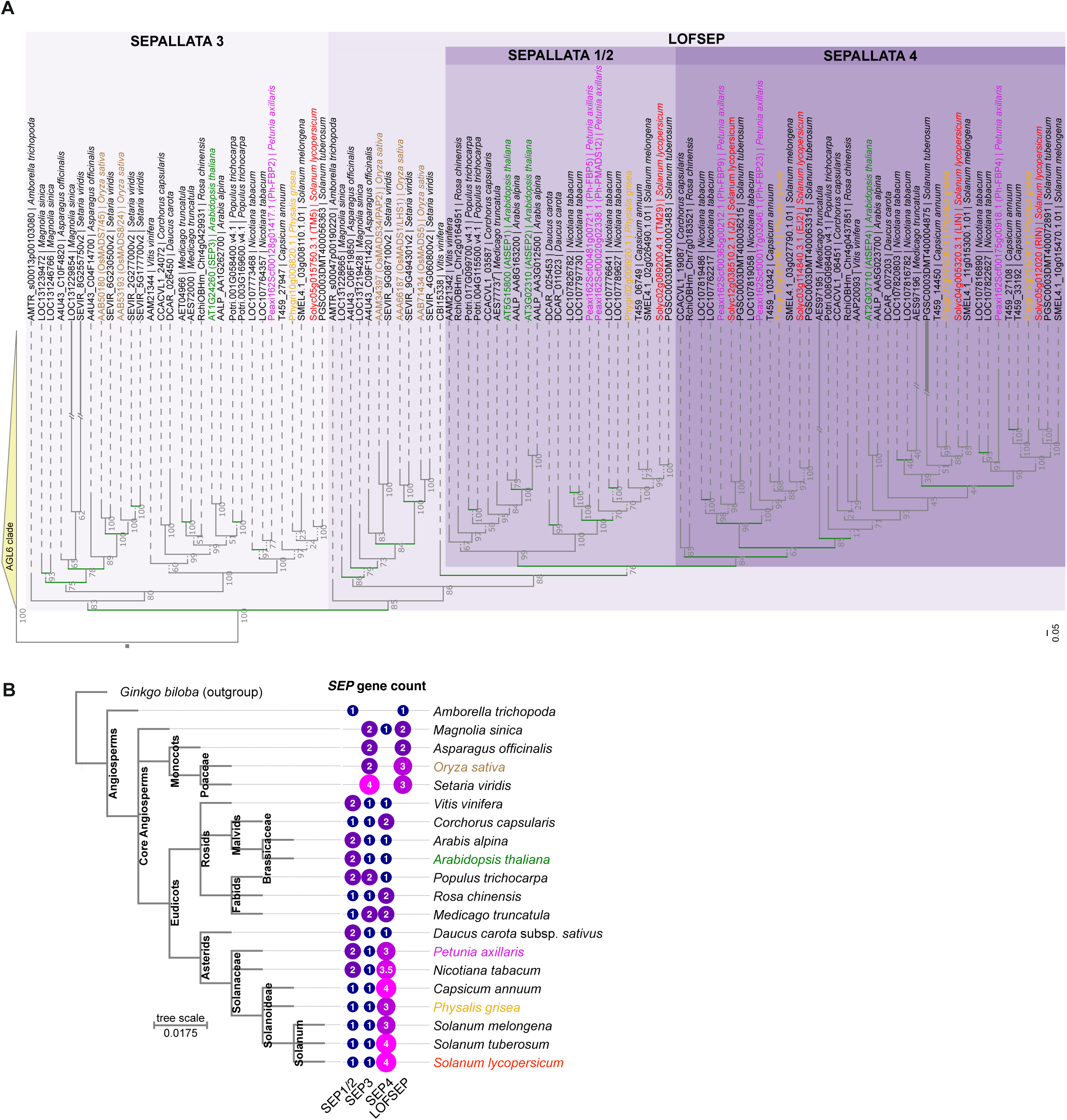
Phylogenetic analysis of the SEPALLATA MADS-box proteins across angiosperms. **(A)** Maximum-likelihood tree of SEP proteins in different angiosperm species. SEP1/2, SEP3, SEP4, and LOFSEP subclades are highlighted in different shades of purple, and the AGL6 sister clade as collapsed outgroup in yellow. Tomato and Arabidopsis genes are labelled in red and green font, respectively. Numbers at nodes represent bootstrap support values from 1,000 replicates. Branch lengths are proportional to the number of amino acid substitutions and are displayed for tomato and Arabidopsis proteins at the branch end points. Branches in green represent duplication events. The scale indicates the average number of substitutions per site. **(B)** Species tree with SEP gene counts across distinct subclades for angiosperm species shown in the phylogenetic tree in (A). Note that gene content of tetraploid N. tabacum is halved to allow comparisons with the other (diploid) genomes.

**Figure S4:**
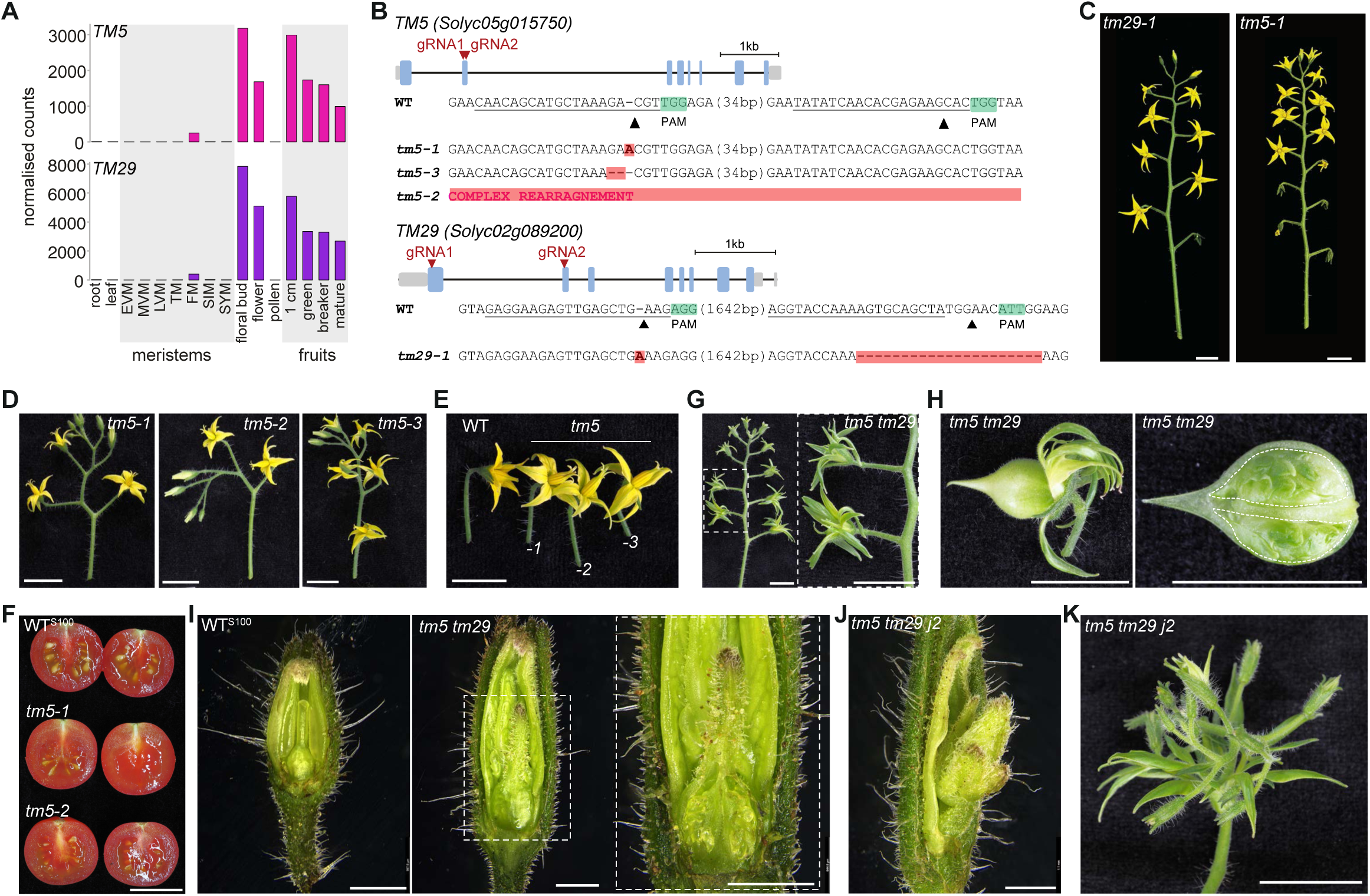
Mutations in the SEPALLATA1/2 paralogs TM5 and TM29 lead to defects in floral organ development. **(A)** Normalized read counts (transcripts per million, TPM) for TM5 and TM29 in different tissues and meristem stages (EVM, early vegetative; MVM, middle vegetative; LVM, late vegetative; TM, transition; FM, floral; SIM, sympodial inflorescence; SYM, sympodial shoot meristem). **(B)** Schematic representation of TM5 (top) and TM29 (bottom) with guide RNAs (gRNA) target positions indicated by arrowheads. Blue boxes represent exons and grey boxes represent UTRs (untranslated regions). The Cas9 cleavage sites for gRNAs are indicated with arrowheads. Mutant allele sequences identified by Sanger sequencing are shown with mutated nucleotide positions highlighted with red background. The Cas9 cleavage sites for guide RNAs are indicated with arrowheads and PAMs are marked in green. PAM, protospacer adjacent motif. **(C)** Photographs of detached inflorescences from the tm29-1 and tm5-1 single mutants. **(D-E)**, Detached inflorescences (D) and flowers (E) from three independent tm5 mutant alleles. Note the outward-bending anthercone in tm5 mutants. **(F)** Fruits of the WT and two independent tm5 mutant alleles. Note the reduced seed number and size in tm5 mutants. **(G)** Image of a representative inflorescence from the tm5 tm29 double mutant with leaf-like floral organs. **(H)** Macroscopic image of a tm5 tm29 flower with a developing pseudofruit (left) and a opened pseudofruit showing two locules filled with leaf-like organs. **(I)** Stereomicroscope images of developing carpels of the WT and tm5 tm29 double mutant. Note that the tm5 tm29 ovary in the image on the right has been opened to expose vegetative organs. **(J)** Stereomicroscope image of an tm5 tm29 j2 triple mutant flower. Note the emerging inflorescence with flower buds at the center of the flower. **(K)** Macroscopic image of a tm5 tm29 j2 flower. Note the vegetative outer organs and the inflorescence in the center of the flower. Scale bars: 1 cm in (C-H), (K) and 1 mm in (I-J).

**Figure S5:**
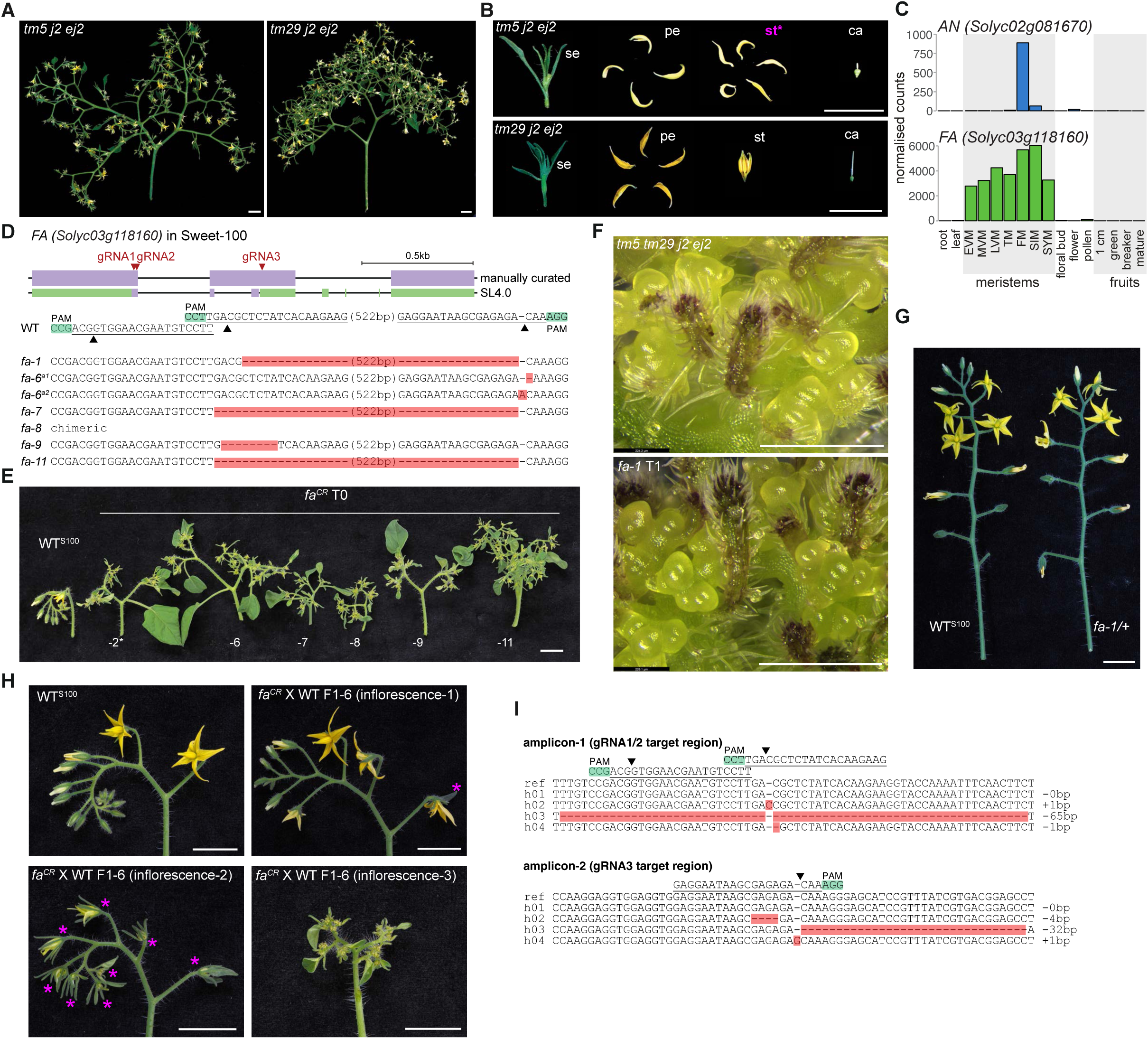
Reductions of SEP1/2/3/4 activity and FALSIFLORA lead to floral organ defects and meristem overproliferation. **(A-B)** Images of tm5 j2 ej2 and tm29 j2 ej2 inflorescences (A) and dissected flowers (B). Se, sepals; pe, petals, st, stamen, ca, carpel. **(C)** Normalized read counts (transcripts per million, TPM) for AN and FA in different tissues and meristem stages (EVM, early vegetative; MVM, middle vegetative; LVM, late vegetative; TM, transition; FM, floral; SIM, sympodial inflorescence; SYM, sympodial shoot meristem). **(D)** Schematic representation of FA with guide RNAs (gRNA) target positions indicated by arrowheads. Purple boxes represent exons and green boxes represent UTRs (untranslated regions). Note that the SL4.0 reference annotation has been manually corrected. The Cas9 cleavage sites for gRNAs are indicated with arrowheads. Mutant allele sequences (identified by Sanger) are shown with mutated nucleotide positions highlighted in red background. The Cas9 cleavage sites for guide RNAs are indicated with arrowheads and PAMs are marked in green. PAM, protospacer adjacent motif. Note that fa-6 and fa-8 are biallelic and chimeric plants, respectively. **(E)** Images of inflorescences from the Sweet-100 WT (WTS100) and six independent first-generation (T0) fa-CRISPR (faCR) transgenics. **(F)** Stereomicroscope images of inflorescence tissue from the tm5 tm29 j2 ej2 quadruple mutant and a homozygous fa-1 T1 plant. **(G)** Images of inflorescences of the WT and a fa-1/+ heterozygote. **(H)** Images of inflorescences of the WT and different sectors of a fa^CR^ X WT F1 individual. Floral organ defects are labelled with magenta asterisks. Note the cauliflower inflorescence on the bottom-right. **(I)** Haplotype sequences at gRNA1/2 target (top) and gRNA3 target (bottom) are shown with mutated nucleotide positions highlighted in red background. The Cas9 cleavage sites for guide RNAs are indicated with arrowheads and PAMs are marked in green. PAM, protospacer adjacent motif. Scale bars: 1 cm in (A-B), (E), (G-H) and 500 µm in (F).

**Figure S6:**
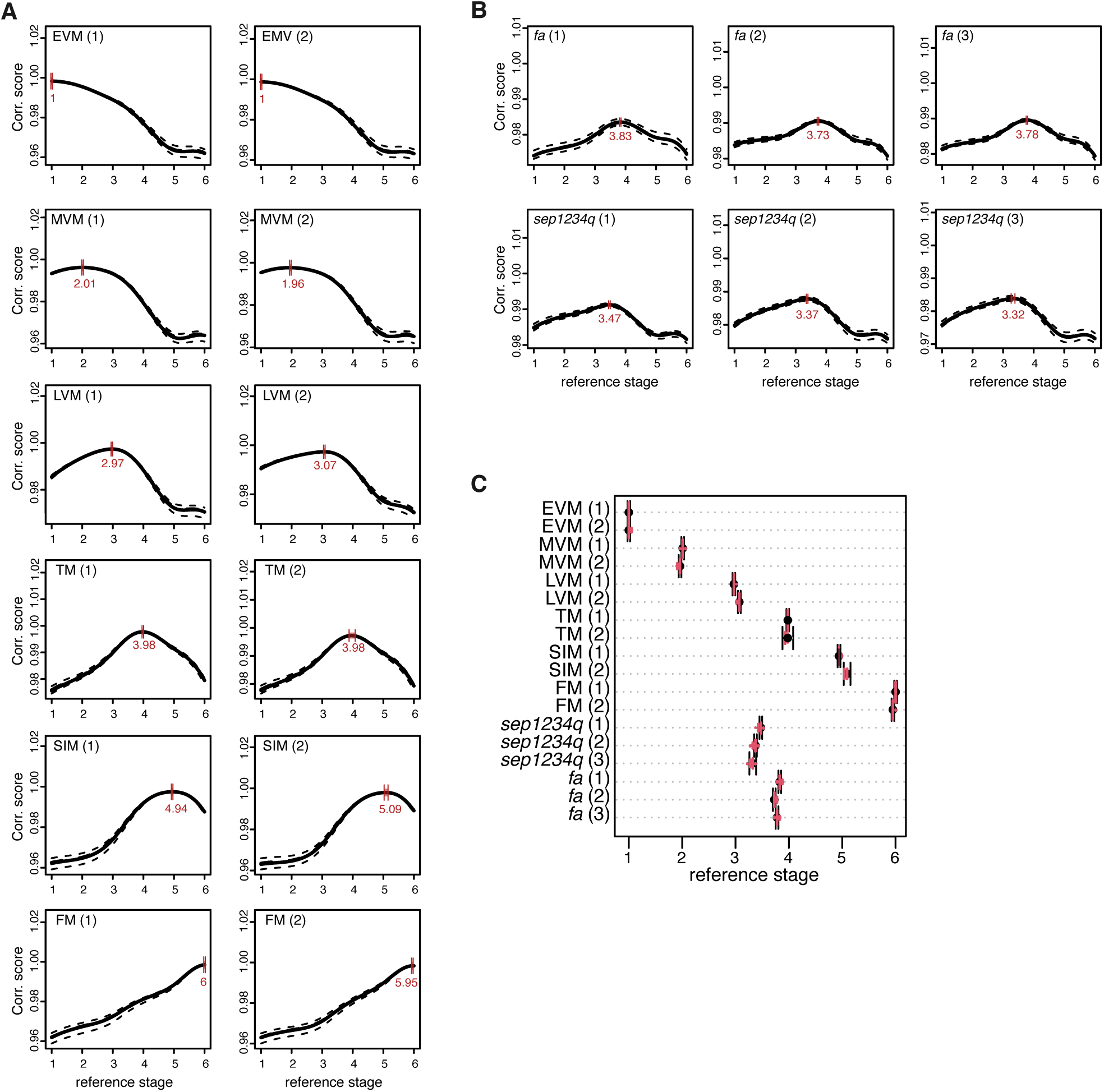
Developmental stage prediction of fa and sep1234q mutant meristem tissue using RAPToR. **(A-B)** Correlation profiles for staged WT meristem (A) and fa and sep1234q mutant meristem (B) samples. Red bars indicate estimation confidence intervals, and black dotted lines indicate the 95% interval of bootstrap correlation with the reference. The sample age estimate is displayed below the interval. Estimated reference age is indicated in red font. **(C)** Developmental stage estimation for staged WT meristem samples and fa and sep1234q mutant meristem. Error bars indicate lower and upper bounds of the estimation confidence intervals, and vertical red lines indicate bootstrap estimates.

